# Sex Specific Attenuation of Reward Preference

**DOI:** 10.1101/2025.02.09.637308

**Authors:** Christopher A. Turner, Jill B. Becker

**Affiliations:** Department of Psychology, University of Michigan; Michigan Neuroscience Institute, University of Michigan

## Abstract

Estradiol receptor signaling has a sex-specific impact on the brain’s reward pathways, enhancing cocaine reinforcement in females but not in males. Selective activation of G-Protein Coupled Estradiol Receptor 1 (GPER-1) in the dorsolateral striatum (DLS) attenuates the reinforcing effects of 0.1% saccharin (SACC) and cocaine in males but not females. This study investigated GPER-1 activation in the DLS and systemically using the GPER-1 agonist, G1, to assess its effect on SACC and cocaine preference in male and female rats. Five experiments were conducted using gonad-intact and gonadectomized animals to determine dose-response effects and the influence of circulating hormones. Intra-DLS GPER-1 activation with 20% G1 selectively reduced SACC preference in intact males but not females, while higher and lower concentrations had no effect. Systemic G1 administration attenuated cocaine-induced conditioned place preference (CPP) in both sexes in a dose-dependent way. Interestingly, systemic administration of G1 did not alter SACC preference in either sex, regardless of the presence or absence of gonadal hormones. These findings suggest that GPER-1 activation influences reward processing in a site, reward, and sex-dependent manner.

**Highlights:**

- Selective activation of the membrane-associated estradiol receptor, GPER-1, in the dorsolateral striatum (DLS) attenuates preference for a 0.1% saccharin solution (SACC) in male but not female rats.
- Systemic activation of the membrane-associated estradiol receptor, GPER-1, attenuates cocaine-induced condition place preference (CPP) in both males and females in a dose-dependent way.
- Systemic GPER-1 activation does not influence SACC preference in either sex.

## Introduction

The sex of an individual is an essential but often overlooked factor when assessing addiction characteristics. Women are known to escalate their drug use earlier, a phenomenon known as “telescoping” (Anglin et al., 1987; Becker and Chartoff, 2019; Griffin, 1989; Haas and Peters, 2000). As a result, despite using drugs for a shorter period, they enter treatment sooner and present with more severe symptoms compared to men (Griffin, 1989; Longshore et al., 1993; McCance-Katz et al., 1999; White et al., 1996). Women also have a longer road to recovery than men. Cocaine withdrawal in women leads to shorter periods of abstinence and higher rates of relapse compared to men (Hudson and Stamp, 2011; Kosten et al., 1993). In addition, following relapse, women use drugs for longer periods than men prior to another attempt at quitting (Gallop et al., 2007; White et al., 1996). These observations indicate that cocaine addiction is more severe in women than in men, suggesting an underlying biological mechanism contributing to these sex differences.

Understanding the neurobiological mechanisms underlying these differences is a critical next step. A prominent mechanism contributing to the sex differences is the sex-dependent actions of gonadal hormones such as estradiol (E2). E2 modulates motivated behaviors such as obtaining food, water, social interaction, or reproduction (Yoest et al., 2014). However, E2 activity can also drive maladaptive behaviors like drug-seeking and -taking behaviors by altering reward processing (Yoest et al., 2014). Following drug use, women reported a greater “high” during the follicular phase of their estrous cycle when E2 levels were at their highest (Evans et al., 2002; Sofuoglu et al., 1999). Animal studies further support this, showing that E2 receptor activity increases the reinforcing effects of drugs of abuse in females (Cummings et al., 2014; Hu and Becker, 2003; Song et al., 2019). E2 receptor signaling causes female rats to acquire drug self-administration and escalate drug use sooner than males (Hu et al., 2004; Lynch et al., 2001). Following E2 activation, female rats also demonstrate increased motivation for drugs, greater cue-reactivity, and more robust reinstatement than males (Becker and Hu, 2008). The effects of E2 on the behavioral response to drugs of abuse highlight the need for further investigation of the interaction between the neurocircuitry underlying motivational processes and E2.

There are three E2 receptor subtypes present in the brain that mediate the effects of E2 activity: E2 receptor alpha (ERα), E2 receptor beta (ERβ), and G-protein coupled estradiol receptor-1 (GPER-1). Most of ERα and ERβ reside in their inactive form in the cytosol. When activated by E2, they dimerize and migrate into the nucleus to modify gene expression (Björnström and Sjöberg, 2005; Nilsson et al., 2001). However, these receptors can also be palmitoylated and trafficked to the plasma membrane by caveolin-1, a structural coat protein. Once at the plasma membrane, they are anchored to plasma membrane invaginations called caveolae. Extranuclear receptors at the plasma membrane, such as metabotropic glutamate receptors, facilitate E2 signaling via protein-kinase cascades (Levin and Hammes, 2016; Lösel and Wehling, 2003; Razandi et al., 2004). GPER-1 embeds itself in various cell membranes depending on the type of neuronal tissue (Brailoiu et al., 2007; Funakoshi et al., 2006; Revankar et al., 2005). Notably, these E2 receptors are expressed in the DLS, a region associated with the development of substance use disorders (Almey et al., 2022, 2016, 2012; Everitt and Robbins, 2013; Krentzel et al., 2021; Quigley and Becker, 2021).

Historically, research has focused on the impact of ERα and ERβ activity as they have confirmed roles altering reward processing in females (Morissette et al., 2008). However, recent studies demonstrate that GPER-1 activity can modulate male reward preference. Knocking out GPER-1 in male mice enhances the development of morphine-induced conditioned place preference (Sun et al., 2020). Studies in rats show that GPER-1 activation attenuates preference formation for substances like saccharin and cocaine in a sex-specific manner (Quigley and Becker, 2021). Specifically, intra-dorsolateral striatum (DLS) administration of G1, a GPER-1 agonist, resulted in a reduced preference for 0.1% saccharin (SACC) vs water. Additionally, it attenuated the development of a cocaine (10 mg/kg) conditioned place preference (CPP) in intact males but not in intact females (Quigley and Becker, 2021). These exciting results suggest that GPER-1-mediated signaling is uniquely positioned to cause behaviorally relevant modifications to male reward processing.

Building on these findings, the experiments presented here aim to characterize further the effects of G1 administration and GPER-1 activation on reward preference in male and female rats. In previous studies, only a single concentration of G1 was used to investigate the effects of DLS GPER-1 signaling on reward preference in intact males and females (Quigley and Becker, 2021). The previous study’s use of a single concentration of G1 may have limited the ability to detect effects in females, who might require a different dose range due to sex-specific pharmacodynamics (Becker, 1990; Tanapat et al., 2005). Using multiple doses and comparing between gonadectomized and intact subjects, the current studies aim to determine dose-response characteristics and effective doses for modulating preference in females. Additionally, we explore the impact of different timings and routes of G1 administration to better understand the role of GPER-1 in reward processing.

## Materials & Methods

### Animals

Male and female Sprague-Dawley rats were obtained from Charles River Breeding Laboratory (Portage, MI) and were approximately 75 days old upon arrival. Animals were maintained in ventilated laboratory cages on a 14:10 light-dark cycle in a temperature-controlled climate of 72 ± 2 °F. Rats had *ad libitum* access to water and phytoestrogen-free rat chow (2017 Teklad Global, 14% protein rodent maintenance diet, Harlan rat chow; Harlan Teklad). Animals were initially housed in same-sex pairs until undergoing surgery, after which they were housed individually. All animals were weighed daily to determine good health, and at this time, females were also vaginally lavaged to determine the stage of estrus. All procedures were performed according to protocols approved by the University of Michigan Institutional Animal Care and Use Committee.

### Stereotaxic Surgery

For studies with craniotomies, animals received bilateral implants of 22-gauge stainless steel guide cannula one week after arrival aimed at the DLS (DLS; AP + 0.4 ML ± 3.6 DV -4.0). On the day of surgery, rats were injected with carprofen (5 mg/kg s.c.) and 30 minutes later were anesthetized with ketamine (50 mg/kg i.p.) and dexmedetomidine (0.25 mg/kg i.p.), then prepared in a stereotaxic frame. After the surgery, rats were given atipamezole hydrochloride (0.5 mg/kg i.p.) and 3 ml 0.9% saline (s.c.). Rats were given carprofen (5 mg/kg s.c.) prophylactically for post-operative pain every 24 hours for three days post-surgery. No animal underwent behavioral testing for at least seven days after surgery.

During surgery, 33-gauge solid stylets were inserted into the 26-gauge hollow guide cannula that was fixed on the animals’ skull. These stylets were flush with the bottom of the guide cannula and did not protrude into the brain. Treatment conditions were randomly assigned to animals prior to behavioral testing. Control animals received 100% cholesterol (CHOL), and experimental animals received either 10, 20, or 30% G1 (agonist targeting GPER1) dissolved in ethanol with cholesterol and then recrystallized. Treatments were delivered via powder packed into stylets, which protruded from the guide cannula by 1 mm and delivered treatment directly into the DLS. Treatment stylets were prepared as previously described (Becker et al., 1987; Quigley and Becker, 2021). When inserting stylets, rats were briefly anesthetized with 5% isoflurane. Data collection began six hours after stylet insertion.

Hollow guides and interlocking treatment stylets were manufactured by and purchased from P1 Technologies (Roanoke, VA). Drugs were obtained from the following sources: G1 (Cayman Chemicals, Ann Arbor, MI; purity ≥ 98%) and cholesterol (Santa Cruz Biotechnology, purity ≥ 92%).

### Gonadectomies

For studies with gonadectomized animals, on the day of surgery, rats were injected with carprofen (5 mg/kg s.c.) and 30 minutes later were placed under isoflurane anesthesia. Females were ovariectomized (OVX), and males were castrated (CAST). Briefly, OVX was performed via a single dorsal incision along the midline below the ribcage. One incision was made on each side laterally through the muscle wall. Each ovary was externalized and then removed following cauterization of the connective tissue. The tissue was returned to the abdominal cavity, and the muscle was sutured on each side. The external incision was closed with an 11 mm wound clip. For CAST, the testes were removed via a ventral approach. A small incision was made along the bottom of the scrotum. The scrotal sac was opened to externalize each testicle and vas deferens before removal. The connective tissue was sutured with surgical thread. The incision site was closed via an 11 mm wound clip. Animals were given one week to recover. Following recovery, vaginal lavage samples were collected from females to confirm the absence of the estrous cycle. Additional details regarding surgeries are published in (Hu and Becker, 2003).

### Osmotic Pump Surgeries

For studies with osmotic pumps, animals received osmotic pump implants while under isoflurane anesthesia. A mid-scapular incision approximately 2-3 cm long was made adjacent to the site chosen for pump placement. A hemostat was inserted into the incision, and a subcutaneous pocket was made large enough to allow free movement of the pump (e.g., 1 cm longer than the pump). The pump containing vehicle [Dimethylsulfoxide (DMSO)] or treatment G1 (90 µg/day in DMSO) was inserted into the pocket, delivery portal first, which minimizes interaction between the compound delivered and the healing of the incision. The pump location was checked to ensure it did not rest immediately beneath the incision. The wound was closed with wound clips or absorbable sutures.

Mini-osmotic pumps were manufactured by and purchased from Alzet (model 2004; Cupertino, CA). Drugs were obtained from the following sources: G1 (Cayman Chemicals, Ann Arbor, MI; purity ≥ 98%); DMSO (Fisher Scientific, Fair Lawn, NJ; purity ≥ 95%)

### Drug Preparation

For subcutaneous injections, G1 was suspended in 0.1% gelatin at 0, 20, or 30 µg/kg concentrations. Solutions were stored at 4°C until the day of use when the syringes were prepared and brought to room temperature. All injections occurred one hour prior to behavioral testing. Hu et al. demonstrated that the effects of E2 treatment are still present one hour after subcutaneous injection (Hu et al., 2004). All hormones and agonists were used within two weeks of preparation.

### Two Bottle Choice

Daily preference of 0.1% SACC versus water was calculated as a percentage using the formula: (0.1 % SACC consumed (g))/(0.1% SACC (g) + water consumed (g))*100. Fluid, SACC, and water intake were normalized to individual body weight, intake (mL)/body weight (kg) before analyses.

The placement of the bottles was switched daily to account for a potential side preference.

#### Experiment 1: Does intra-DLS activation of GPER-1 influence SACC preference and fluid intake in a concentration- and sex-dependent manner?

This experiment investigates the concentration- and sex-dependent effects of targeted GPER-1 activation in the DLS on SACC preference, hypothesizing that GPER-1 activation will alter reward-seeking behavior in a manner consistent with observed sex differences. Two bottles were accessible to individually housed animals in their home cages. On days 1-4, both bottles contained water only. During days 5-12, one bottle contained water, and the other contained a solution of 0.1% SACC (Sigma-Aldrich, St. Louis, MO; purity ≥ 92%) dissolved in water. Treatment stylets (0, 10, 20, or 30% G1 in cholesterol) were inserted into the DLS four hours before SACC introduction on day five and remained for the duration of the experiment. Once daily, one hour before the start of the dark cycle, the bottles were removed, weighed, and refilled.

#### Experiment 2: Does systemic administration of G1 influence the development of a cocaine-induced conditioned place preference?

This experiment examines the effects of systemic activation of GPER-1 on the development of a cocaine-induced CPP, hypothesizing that GPER-1 activation will modulate male, but not female, preference formation. Some minor modifications were made to the CPP method described in (Quigley and Becker, 2021). Animals were randomly assigned to one of three treatment groups: vehicle (0.1% gelatin), 20 µg/kg, or 30 µg/kg of G1 suspended in 0.1% gelatin. During conditioning sessions, one hour prior to the start of a conditioning session, animals received an intra-scapular subcutaneous injection of their designated treatment.

#### Experiment 3: Does systemic activation of GPER-1 influence SACC preference and fluid intake in a dose- and sex-dependent manner?

This experiment investigates the concentration- and sex-dependent effects of systemic GPER-1, hypothesizing that GPER-1 activation will alter reward-seeking behavior in a manner consistent with observed sex differences from Experiment 1. Two bottles were accessible to individually housed animals in their home cages. On days 1-4, both bottles contained water only. During days 5-12, one bottle contained water, and the other contained a 0.1% SACC solution. Beginning on day 5, animals received daily subcutaneous injections of G1 (0, 20, 30 µg/kg) suspended in 0.1% gelatin for the duration of the experiment. These doses were selected to match those used in the CPP study. At the time of the injections, bottles were removed, weighed, and refilled.

#### Experiment 4: Does systemic administration of G1 rapidly influence SACC preference in a dose- and sex-dependent manner?

As a membrane-associated receptor, GPER-1 is believed to mediate some of the rapid effects of E2 signaling (Alexander et al., 2017; Revankar et al., 2005). G1 has a half-life of approximately 4 hours in mice (Natale et al., 2020). In Experiment 2, our GPER-1 agonist was administered one hour prior to the conditioning session, during a time window of significant drug availability. Because Experiment 3 examined SACC preference over a 24-hour period, any acute effects of systemic administration of G1 may masked.

Here, we tested whether systemic administration of G1 rapidly influences SACC preference within the drug’s half-life window. We hypothesized that GPER-1 will rapidly alter reward-seeking behavior in a sex specific manner consistent with the results of Experiment 2. Food and water were removed one hour before the lights went off. At this time, rats also received subcutaneous injections of G1 (0, 20, or 30 µg/kg) suspended in 0.1% gelatin. One hour after the injections, rats were given one hour of access to two bottles: one containing water and the other containing a 0.1% SACC solution. After one hour, SACC bottles were removed from the cages; chow and water were returned. Intake was measured by weighing the bottles at the start and finish of the testing period. Animals underwent five consecutive days of testing.

#### Experiment 5: Does chronic systemic administration of G1 influence SACC preference in a sex-dependent manner?

The studies that found an effect of G1 administration on SACC preference had two common points: 1. G1 administration was continuous, and 2. the animals tested were intact. To best replicate those conditions, the final SACC preference study used mini-osmotic pumps to achieve continuous administration of G1 (90 µg/day), and the animals were left intact in case normally circulating hormones were necessary to see an effect. Two bottles were accessible to individually housed animals in their home cages. On days 1-4, both bottles contained water only. During days 5-12, one bottle contained water, and the other contained a solution of 0.1% SACC. Once daily, one hour before the start of the dark cycle, the bottles were removed, weighed, and refilled.

### Euthanasia and Tissue Preparation

Animals received 0.5 ml of Sodium Pentobarbital (i.p). Once the animal was fully sedated, it was perfused transcardially with 0.1M phosphate-buffered saline followed by 4% paraformaldehyde. Each rat’s brain was dissected and post-fixed in 4% paraformaldehyde for 24 hours and stored in 10% sucrose afterward. Brains were sliced on a cryostat in 60-micron sections, mounted on slides, stained with cresyl violet, and cover slipped. Sections were analyzed for accurate guide cannulae placements.

### Statistical Analyses

One-way ANOVA analyses were used to assess treatment differences in mean SACC preference, fluid intake, SACC intake, and water intake between treatments within sex. Tukey post hoc comparisons were performed when significant main effects were detected. If data violated the assumption of equal variance, a Welch ANOVA test was conducted, and Dunnett’s T3 post hoc comparisons were performed when significant main effects were detected. If data violated the assumption of normality, a Kruskal-Wallis test was conducted, and Dunn’s post hoc comparisons were performed when significant main effects were detected.

Two-way repeated measures (RM) mixed models were used to analyze SACC preference, fluid, SACC, and water intake were analyzed within sex with treatments and time as factors.

Two-way ANOVA analyses were used to assess differences in mean SACC preference, fluid intake, SACC intake, and water intake with sex and treatment group as factors. Tukey post hoc comparisons were performed when significant main effects or interactions were detected.

CPP data were analyzed using a RM two-way ANOVA between treatment groups and within sex. A Holm-Sidak post hoc test was used to determine if there were sex differences within each test session. For males and females independently, data were analyzed by time spent in the drug-paired chamber (pre-test vs. test) between treatment conditions within each sex.

All data are presented as means ± SEM. Outliers were identified as values outside of the range of 1.5 IQR for a given sex x treatment x day combination and excluded from analyses.

Two bottle choice and CPP statistical analyses were performed using GraphPad Prism v10.2.3. For effects described as significant, *p* < 0.05 and trends are reported for *p* < 0.06.

## Results

Experiment 1: Intra-DLS G1 administration modulates 0.1% SACC preference in a sex-dependent manner

### SACC Preference

Male preference scores across different G1 treatment groups are shown in Figure 1 (A&B). There were significant group differences between mean preference score (Fig. 1A; F_(3,16)_ = 12.440, 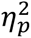 = 0.700). 20% G1 treated males showed a lower mean preference score than 0% (*d =* 2.915), 10% (*d =* 2.060) and 30% (*d = 1*.890) G1 treatment groups. A concentration of 20% G1 administered into the DLS, attenuated SACC reward preference in males compared to all other groups.

**Figure 1.**
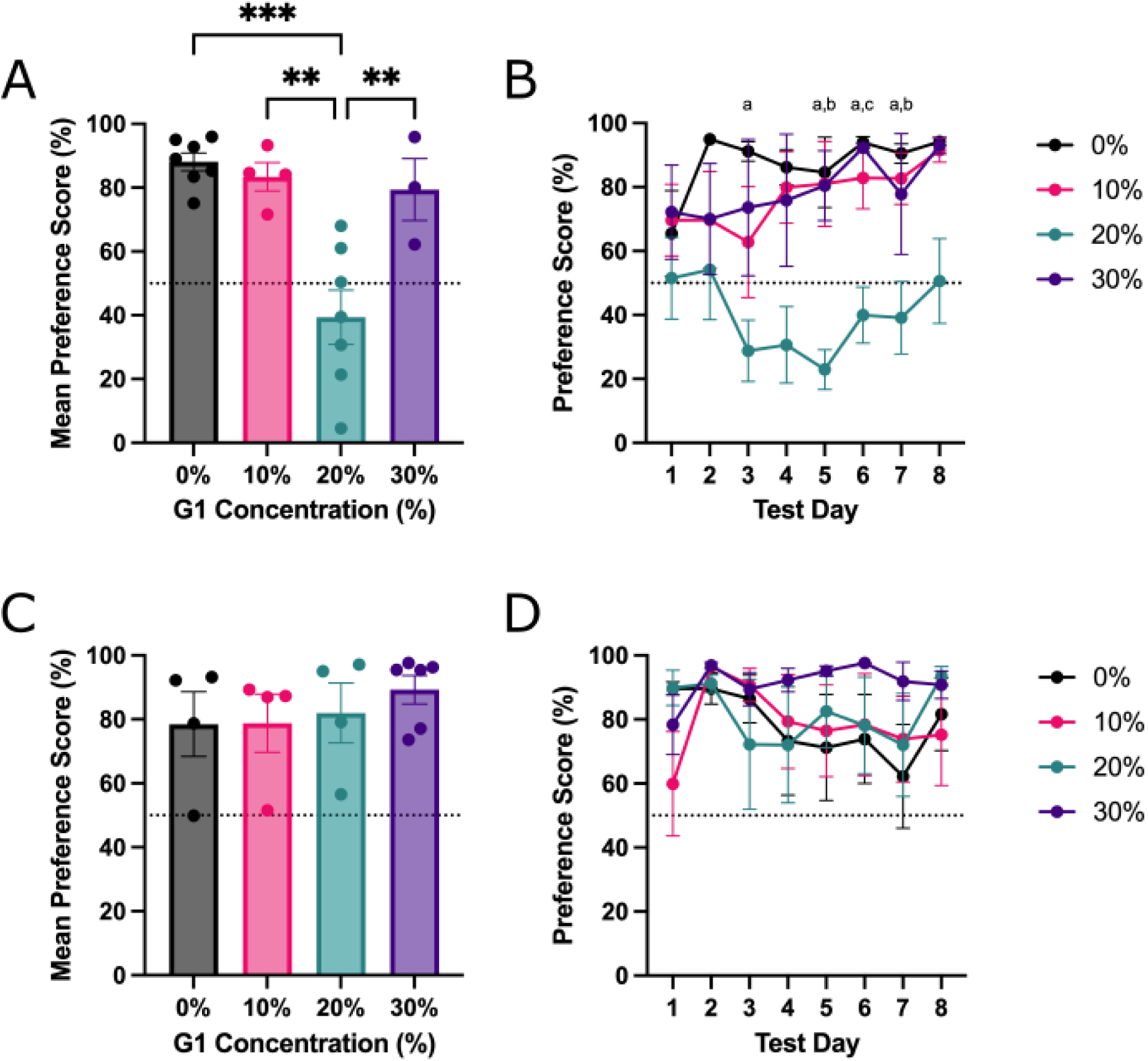
Experiment 1: Intra-DLS GPER-1 activation with 20% G1 attenuates SACC preference in males but not females. (A&B) Males (C&D) Females. (A) 20% G1 males have a significantly lower preference for SACC than all other groups. Males developed a conditioned aversion, as indicated by a mean preference score < 50%. (B) 20% G1 males had significantly lower preference scores than the 0% G1 group on days 5 and lower scores than the 0% and 10% G1 groups on day 7. (C&D) All female treatment groups formed a SACC preference as indicated by a mean preference > 50%. (A&C) Data of individual animals are shown as mean ± SEM. (B&D) Group averages are shown as mean ± SEM. *Note. a: 0% significantly different from 20%, b: 10% significantly different from 20%, c: 30% significantly different from 20%. **p < 0.01, ***p < 0.001*.

As hypothesized, there were significant differences in preference scores between groups across the eight days (Fig. 1B; Effect of Treatment: F_(3,16)_ = 11.110, 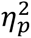 = 0.676). 20% G1-treated males had lower preference scores than 0% G1-treated males on days four through seven (*d* = 2.261, 3.067, 3.354, 2.334) and the 30% G1-treated males on day six (*d* = 2.888). Notably, the effect of GPER-1 activation at this 20% concentration was not due to the duration of the treatment as there was no effect of time or time x treatment interaction.

Figure 1 (C&D) shows female preference across different G1 treatment groups. All treatment groups showed a SACC preference, consistent with previously reported findings (Quigley and Becker, 2021). No effect of treatment, time, or treatment x time interaction was observed for preference score across the eight days. Intra-DLS GPER-1 activation did not attenuate SACC preference in females at any concentration.

Figure 2 shows male and female preference scores. G1 treatment had a significant effect on SACC preference (Fig. 2; Effect of Treatment: F(3, 30) = 4.529; η= 0.312). As expected, the effect of G1 treatment on SACC preference differed between males and females. Specifically, males in the 20% G1-treated males had lower preference scores than 20% G1-treated females (*d* = 2.000).

**Figure 2.**
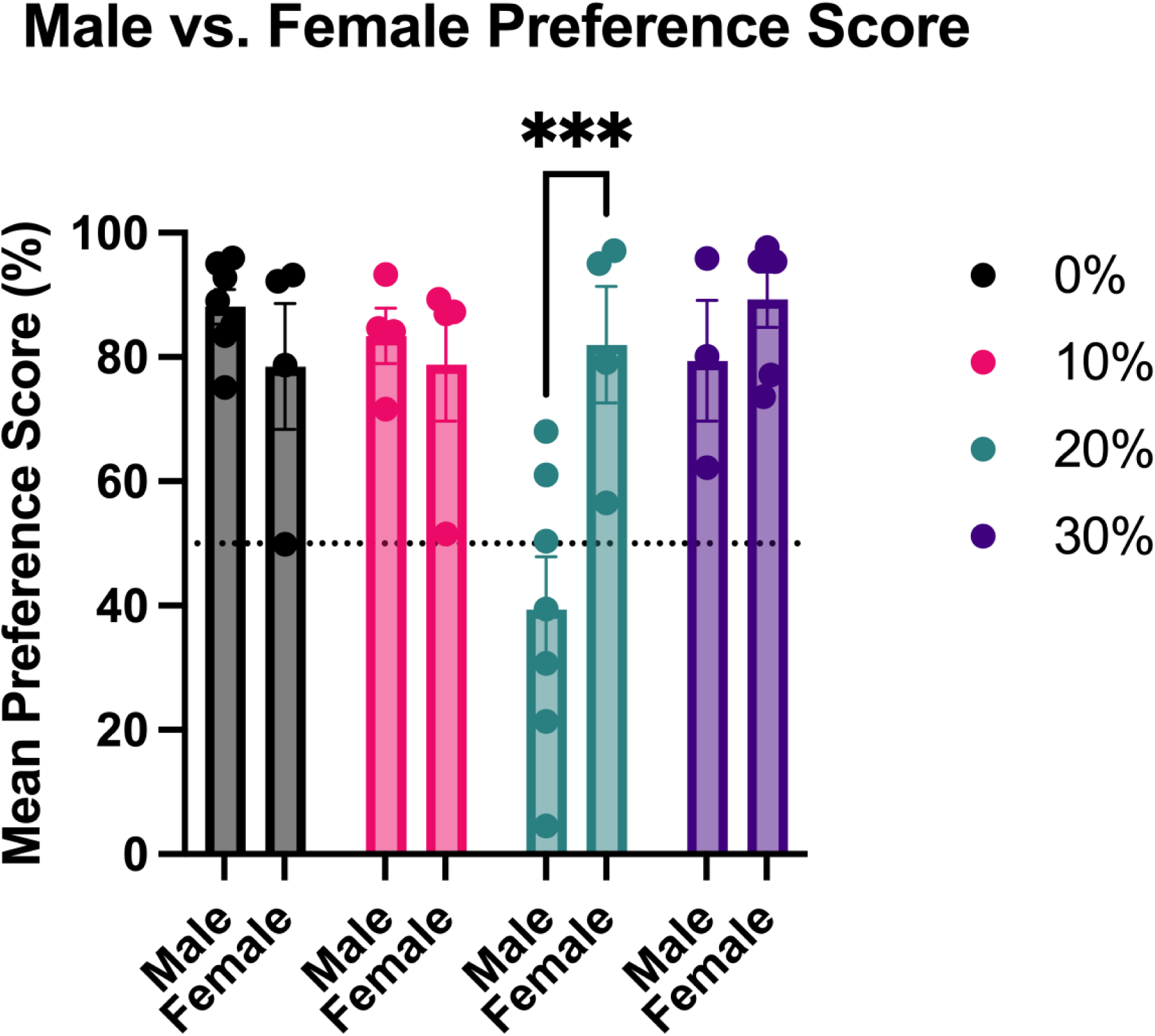
**Experiment 1: Intra-DLS GPER-1 activation attenuates SACC preference in males, but not females**. Intra-DLS activation using 20% G1 significantly reduces SACC preference in males vs. females (Effect of Sex & Sex x Treatment interaction: *p < 0.01*). Data from individual animals are shown as mean ± SEM. ***p < 0.001

### Fluid Intake

Male and female fluid intake values are shown in Figure 3, and M vs. F mean fluid intake values are shown in Figure 4. There were no group differences between treatments within sex or between days within sex (Fig. 3). However, females consumed significantly more fluid compared to males (Fig. 4; Effect of Sex: F_(1, 30)_ = 12.310, 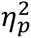 = 0.291). Males in the 20% G1 group drank less fluid overall than females in the 20% G1 group (*d* = 1.350), as did males compared to females in the 30% treatment group (*d* = 1.652).

**Figure 3.**
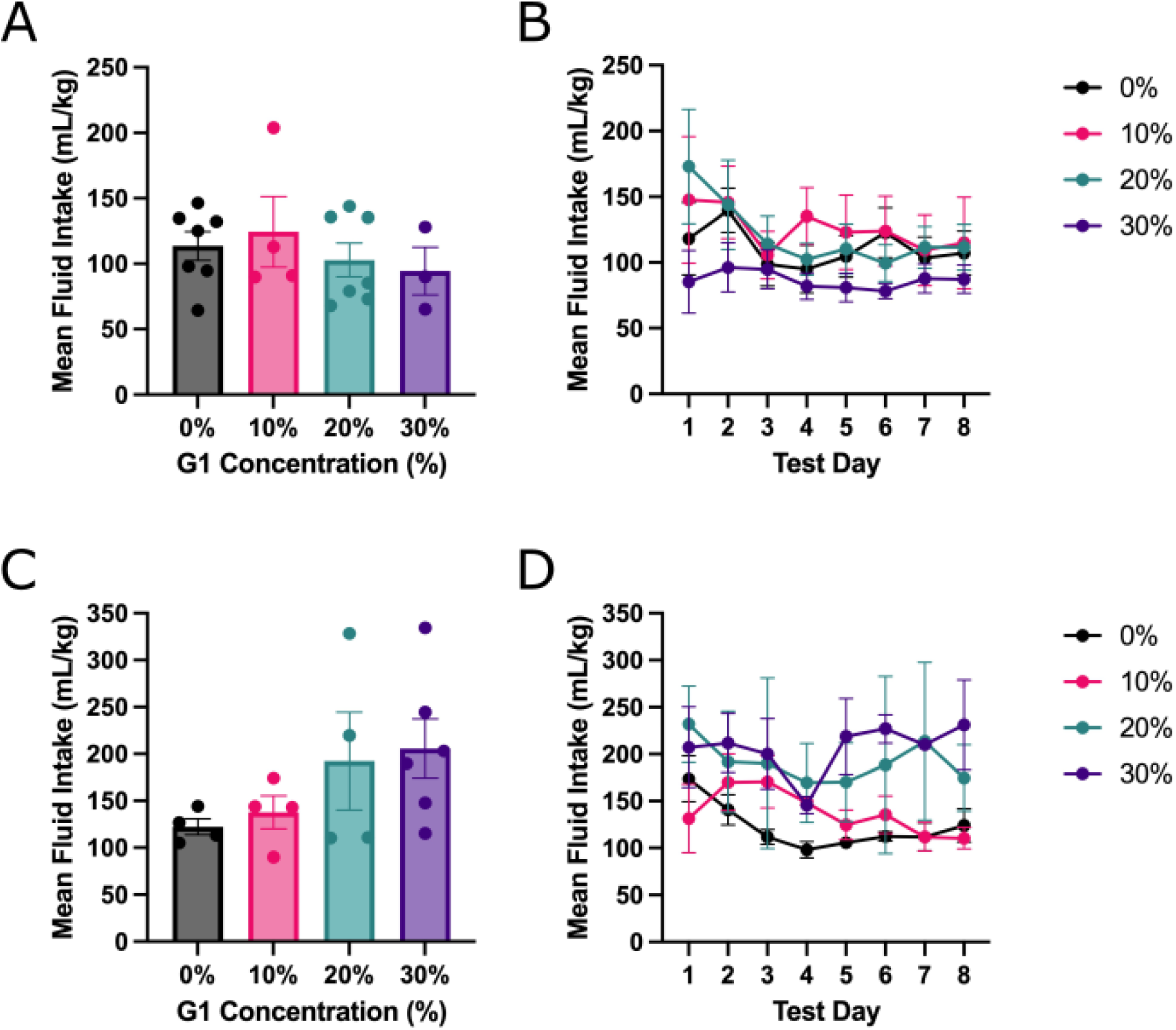
**Experiment 1: Intra-DLS GPER-1 activation did not impact fluid intake in males or females in any treatment group**. (A&B) Males (C&D) Females. All groups showed similar patterns of fluid intake. (A&C) Data of individual animals are shown as mean ± SEM. (B&D) Group averages are shown as mean ± SEM.

**Figure 4.**
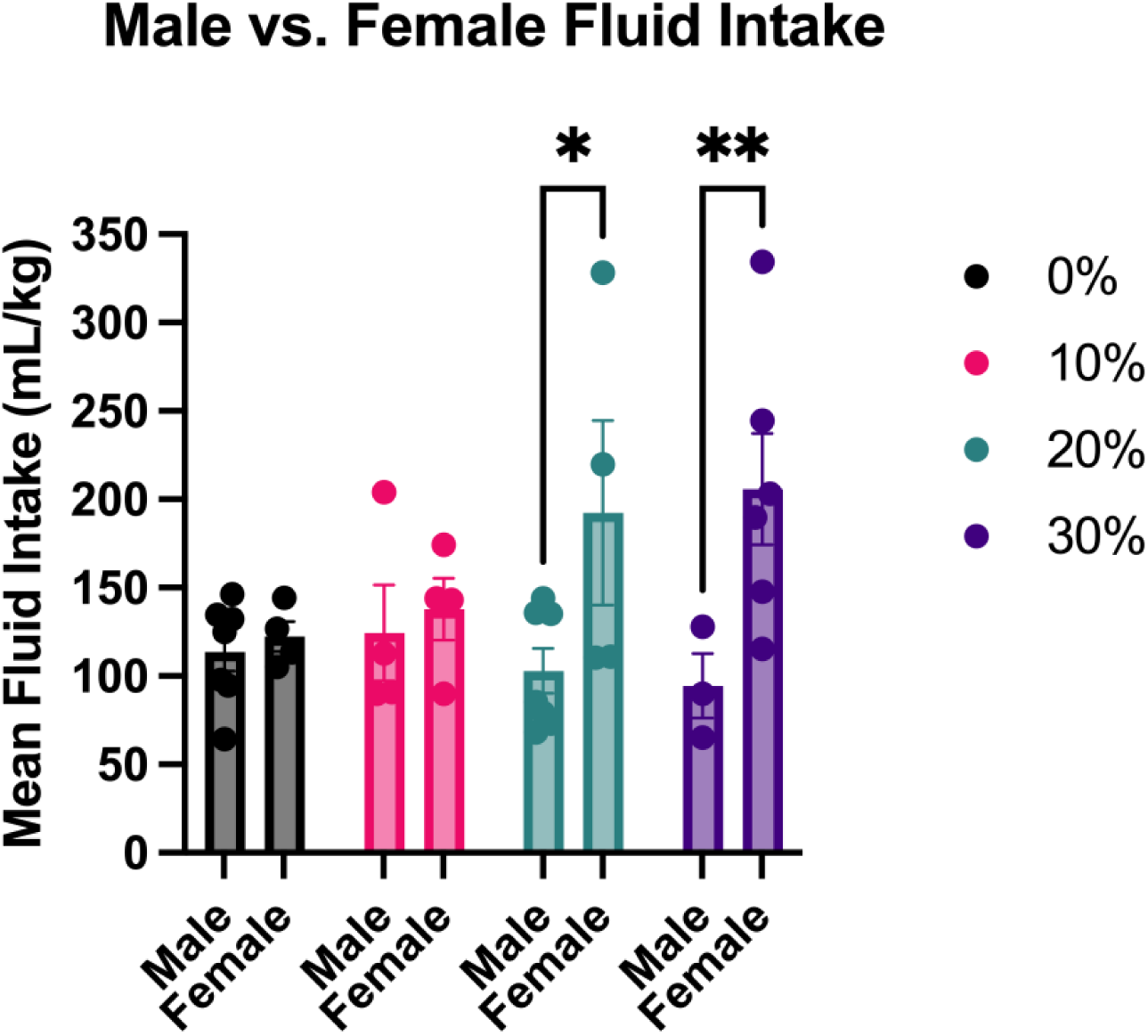
Experiment 1: Intra-DLS GPER-1 activation modulates fluid intake in a sex-dependent manner. Females in both the 20% and 30% G1 groups consumed significantly more fluid than males in the same group (Effect of Sex: *p < 0.01*). Data from individual animals are shown as mean ± SEM. *p < 0.05, **p < 0.01.

### SACC Intake

Male and female mean SACC intake values are shown in Table 1. In males, there was a trend toward a significant difference in mean SACC intake (Table 1; F_(3,16)_ = 0.720, 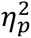 = 0.119). There was a trend toward a significant difference in SACC intake across days (Table 1; Effect of Treatment: F_(3, 16)_ = 3.107, 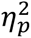 = 0.368), but no effect of Time or Time x Treatment interaction. Mean SACC intake in 0% G1-treated males was significantly greater than in 20% G1-treated males (*d* = 1.610). Female treatment groups did not differ in mean SACC intake or SACC intake across days.

As anticipated, there were group differences between males and females (Table 1; Effect of Sex: F_(1, 30)_ = 10.460, 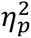 = 0.511). There was no effect of Treatment. Additionally, there was a trend toward a Sex x Treatment interaction (Table 1; F_(3, 30)_ = 2.710, 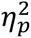 = 0.213). Specifically, males in the 20% G1 treatment group drank less SACC than females in the same group (*d* =1.660). There was also a trend toward 30% G1 males drinking less SACC than 30% G1 females (*d* =1.468).

### Water Intake

Male and female mean water intake values are shown in Table 1. Water intake significantly differed between treatment groups (W_(3.000, 6.422)_ = 17.80, *p* = .002). Specifically, 20% G1 treated males’ water intake was greater than that of the 0%, 10%, and 30% G1 groups (*d* = 4.225, 3.127, and *d* = 3.03, respectively). GPER-1 activation had a significant effect on male water intake over time (Table 2; Effect of Treatment: F_(3, 16)_ = 27.900, 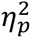 = 0.840). Specifically, males in the 20% G1 group drank more water on day five compared to day one (Table 2; *d* = 1.271) and less water on day six compared to day five (Table 2; *d* = 0.954). In females there were no differences between treatment groups (Data Not Shown).

**Table 1.** Statistical analyses for Experiment 1: Mean SACC and Water Intake. Table 1 shows data from a two-way ANOVA analysis of male and female mean SACC & water intake. There was a significant main effect of Sex for mean SACC intake and a trend toward a Sex x Treatment interaction. Males in the 20% and 30% G1 treatment groups drank significantly less SACC than females in the same group. There was significant main effect of Treatment as well as a Sex x Treatment interaction for mean water intake. Males in the 20% G1 group drank significantly more water than all other male treatment groups. Additionally, 20% G1 males drank more water than 20% G1 females. *Note. S = sex; T = Treatment. Means with different superscripts significantly differ (p < 0.05) as determined by Tukey post hoc tests. *p < 0.05, **p < 0.01*.

| Experiment 1: Mean SACC & Water Intake |  |  |  |  |  |  |  |  |  |  |  |  |
| --- | --- | --- | --- | --- | --- | --- | --- | --- | --- | --- | --- | --- |
| Variable | 0% G1 |  | 10% G1 |  | 20% G1 |  | 30% G1 |  | Two-Way ANOVA |  |  |  |
| | M | SD | M | SD | M | SD | M | SD | Effect | F | dF | $\eta_p^2$ |
| SACC Intake |  |  |  |  |  |  |  |  | S x T | 2.710 | 3,30 | 0.213 |
| Male | 103.751 | 29.578 | 82.617 | 13.874 | 48.807<sup>a</sup> | 38.196 | 80.170<sup>b</sup> | 40.440 | S | 10.460** | 1,30 | 0.259 |
| Female | 98.663 | 24.996 | 111.270 | 39.820 | 171.680<sup>a</sup> | 116.312 | 189.915<sup>b</sup> | 84.699 | T | 0.691 | 3,30 | 0.065 |
| Water Intake |  |  |  |  |  |  |  |  | S x T | 5.370** | 3,30 | 0.349 |
| Male | 9.856<sup>c</sup> | 3.826 | 15.263<sup>d</sup> | 1.773 | 54.101<sup>c,d,e,f</sup> | 14.309 | 14.205<sup>e</sup> | 8.861 | S | 0.111* | 1,30 | 0.004 |
| Female | 23.683 | 20.827 | 26.600 | 27.305 | 20.618<sup>f</sup> | 15.099 | 15.867 | 12.511 | T | 4.671 | 3,30 | 0.318 |

**Table 2.**
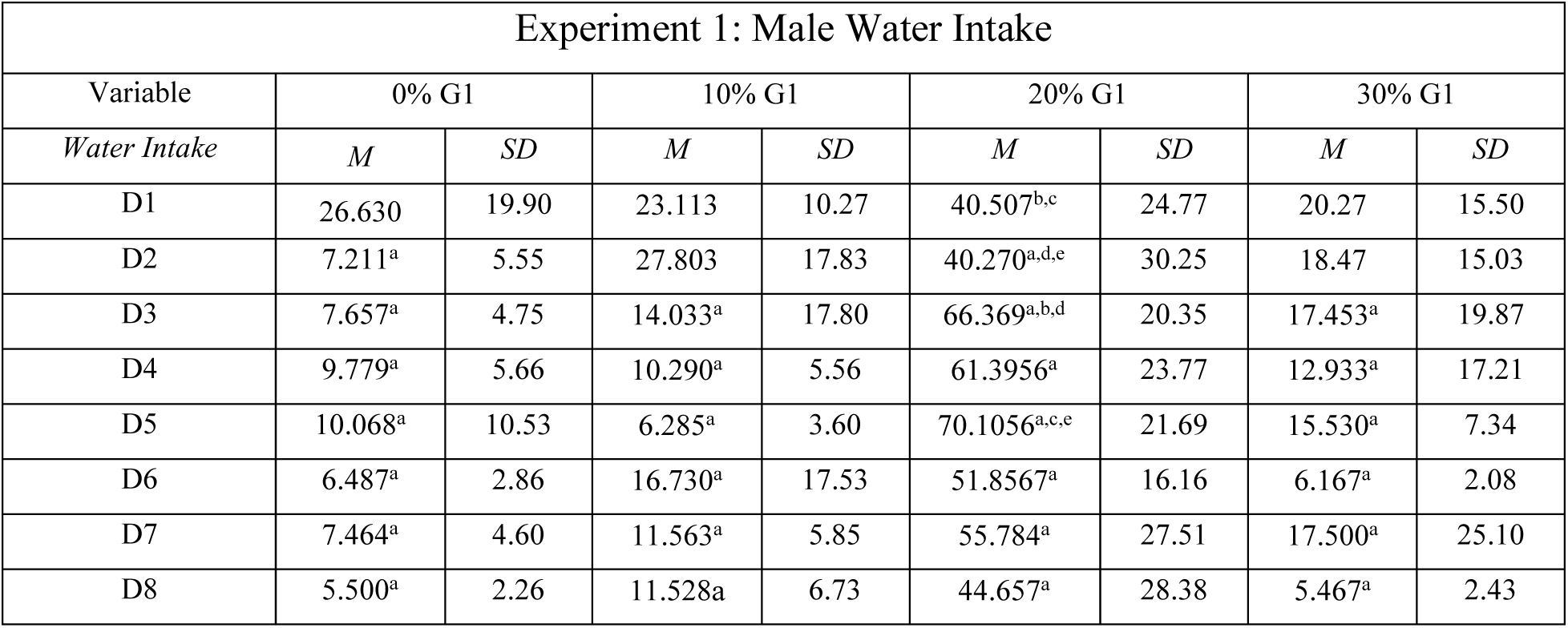
Statistical analyses for Experiment 1: Male Water Intake. Table shows data from Tukey post hoc comparisons of male water intake. *Means with a superscript were significantly different (p < 0.05) as determined by Tukey post hoc tests*.

Mean water intake between males and females differed as a function of both treatment (Table 1; Effect of Treatment: F_(3, 30)_ = 4.671, 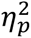 = 0.318) as well as a Sex x Treatment interaction (Table 1; Sex x Treatment Interaction: F_(3, 30)_ = 5.370, 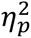 = 0.349. Males in the 20% G1 treatment group drank significantly more water than females in the 20% G1 treatment group (Table 1; *d* = 2.297).

### Summary

Intra-DLS administration of 20% G1 significantly reduced SACC preference in males compared to other treatment groups, with a corresponding decrease in total fluid and SACC intake. Females showed no significant changes in preference or intake.

Experiment 2: Acute systemic administration of G1 attenuates the formation of a cocaine-induced conditioned place preference in males and females

Data on the time spent in the drug-paired chamber (pre-test vs. test) from the control male and female groups were analyzed. There were no sex differences between groups (Fig. 5; Effect of Sex: F_(1,9)_ = 1.369, *p* = 0.272; Sex x Session interaction: F_(1,9)_ = 0.003; *p* = 0.959). Male and female control groups were combined, to increase statistical power.

**Figure 5.**
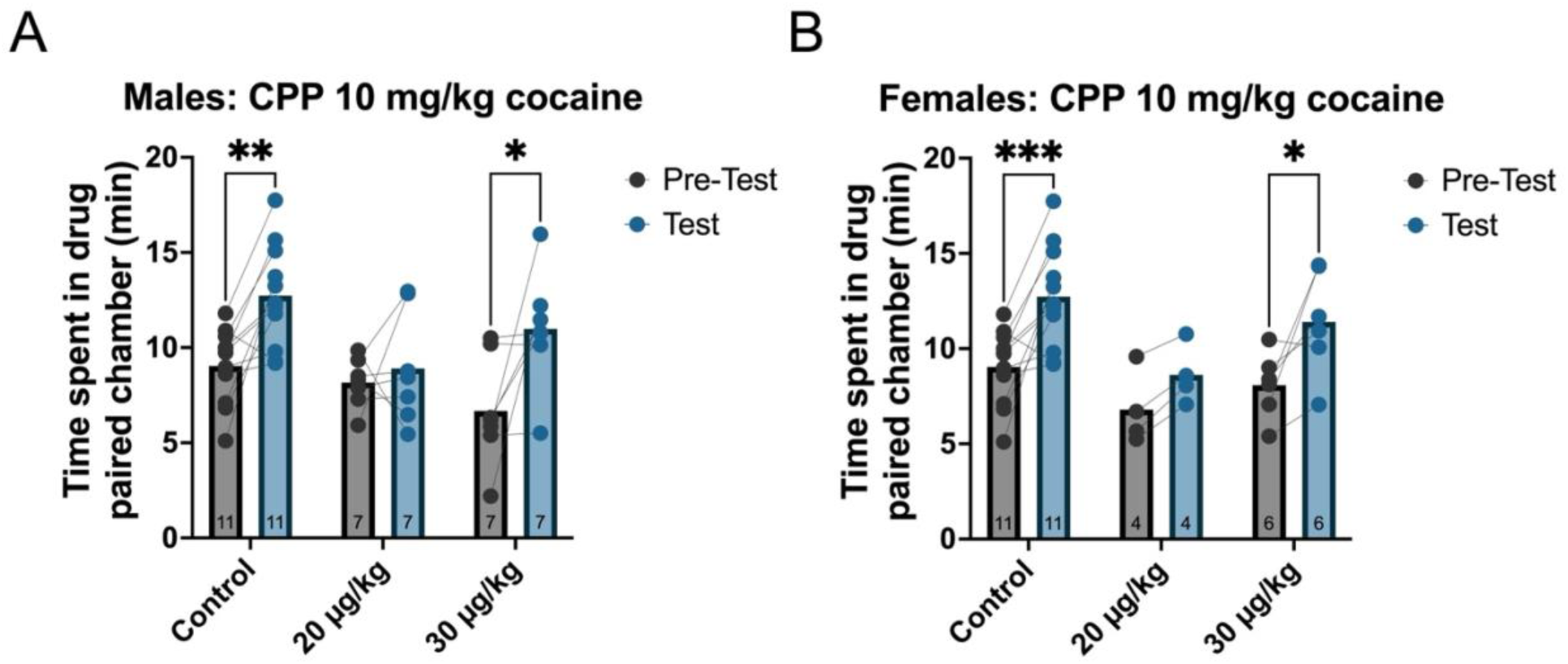
Systemic administration of the GPER-1 agonist, G1, attenuates cocaine CPP in males and females. Activation of GPER-1 via administration of G1 attenuates cocaine CPP in both males and females. (A) At a conditioning dose of 10 mg/kg cocaine, the Control (0.1% gelatin) and 30 µg/kg G1 groups of males developed a cocaine-induced CPP. The 20 µg/kg doses blocked the development of a cocaine CPP. (B) At a conditioning dose of 10 mg/kg cocaine, both the Control (0.1% gelatin) and the 30 µg/kg G1 female treatment groups developed a cocaine-induced CPP. The 20 µg/kg dose attenuated the development of a cocaine CPP. *p < 0.05, **p < 0.01

In males and females, the development of a cocaine-induced conditioned place preference was affected by both test session (Males: F_(1,22)_ = 15.290, 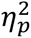 = 0.420; Females: F_(1,18)_ = 20.44, 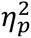 = 0.532) and treatment (Males: F_(2,22)_ = 4.811, 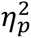 = 0.304; Females: F_(2,18)_ = 4.531; 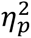 = 0.335). There was no Sex x Session interaction. 20 µg/kg & treated males did not develop a cocaine-induced CPP (*d* = 0.328). Interestingly, 20 µg/kg females also failed to develop a cocaine-induced CPP (*d* = 1.030). All other groups significantly increased their time spent in the cocaine-paired chamber.

Experiment 3: Systemic administration of G1 does not modulate 0.1% SACC preference in gonadectomized animals of either sex

### SACC Preference

CAST male and OVX female preference scores are shown in Figure 6. Within sex, CAST males and OVX females showed a similar mean preference for SACC across all treatment groups. However, in both sexes, preference score did fluctuate across days (Fig. 6; Males – effect of time: F_(4.210, 52.92)_ = 5.507, 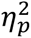 = 0.305; Females - effect of time: F_(3.148, 30.58)_ = 4.490, 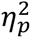 = 0.316). CAST males in the 20 µg/kg group had a mean preference score that was lower on day six compared to day one (*d* = 2.177) or day five (*d* = 2.023). The mean preference score for OVX females in the 0 µg/kg group was significantly lower on day five compared to day eight (*d =* 4.307).

**Figure 6.**
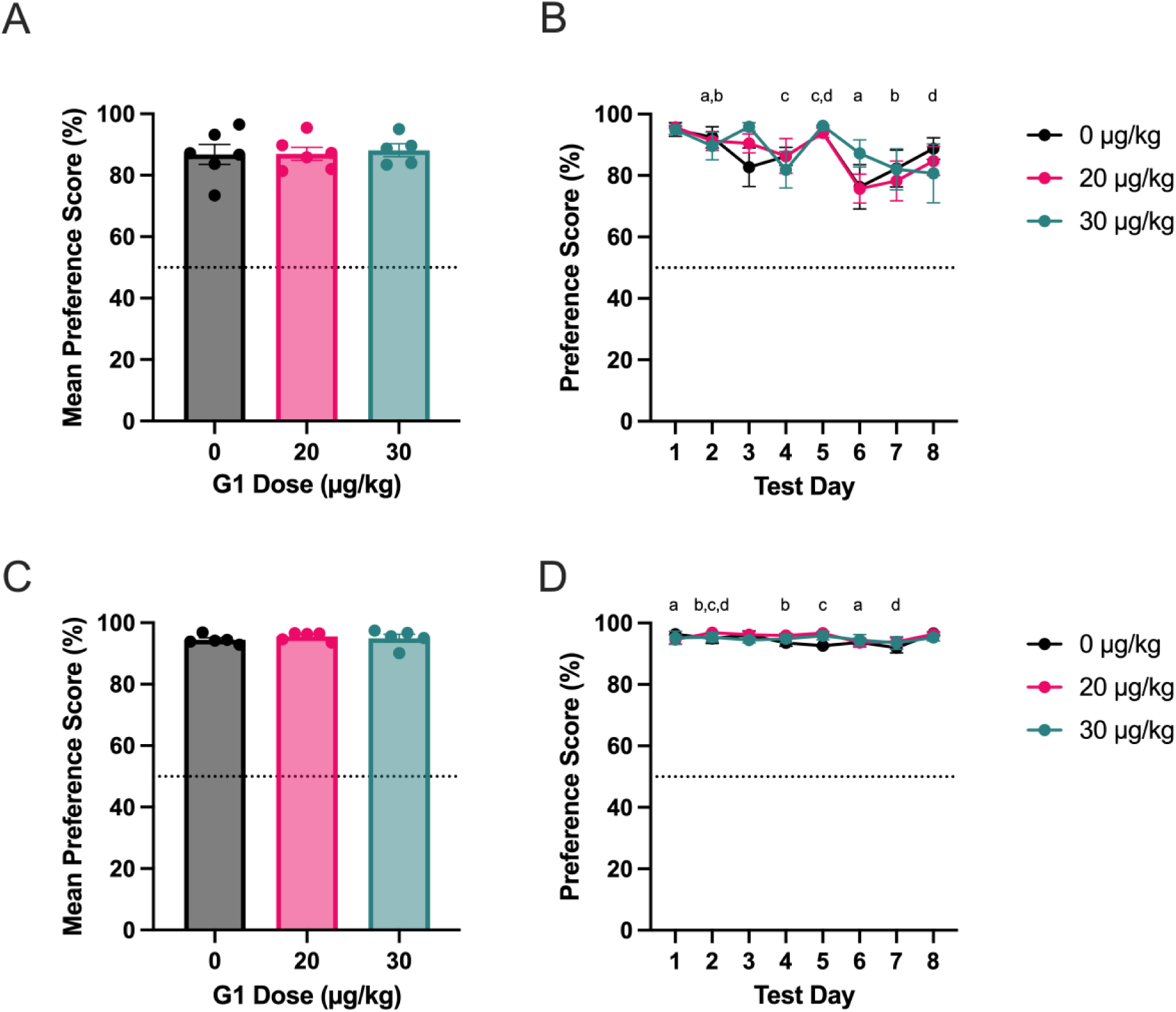
Experiment 3: Systemic administration of G1 does not attenuate preference score in gonadectomized males or females. (A&B) Males (C&D) Females. All treatment groups formed a SACC preference, as indicated by a mean preference score > 50%. Preference score fluctuated across days in both sexes (Effect of Time: *p* < .01) (A&C) Data of individual animals are shown as mean ± SEM. (B&D) Group averages are shown as mean ± SEM. *Note. Days with different letters differ significantly (p < 0.05) as determined by Tukey post hoc tests*.

Figure 7 shows CAST male vs. OVX female mean preference scores. Overall, OVX females showed a greater mean SACC preference than CAST males (Fig. 7; Effect of Sex: F_(1, 27)_ = 20.36, 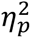 = 0.408). CAST males in 0% and 20% G1 treatment groups had a lower mean preference score than OVX females in the same treatment group (Fig. 7; *d*’s = 1.378, 2.213)

**Figure 7.**
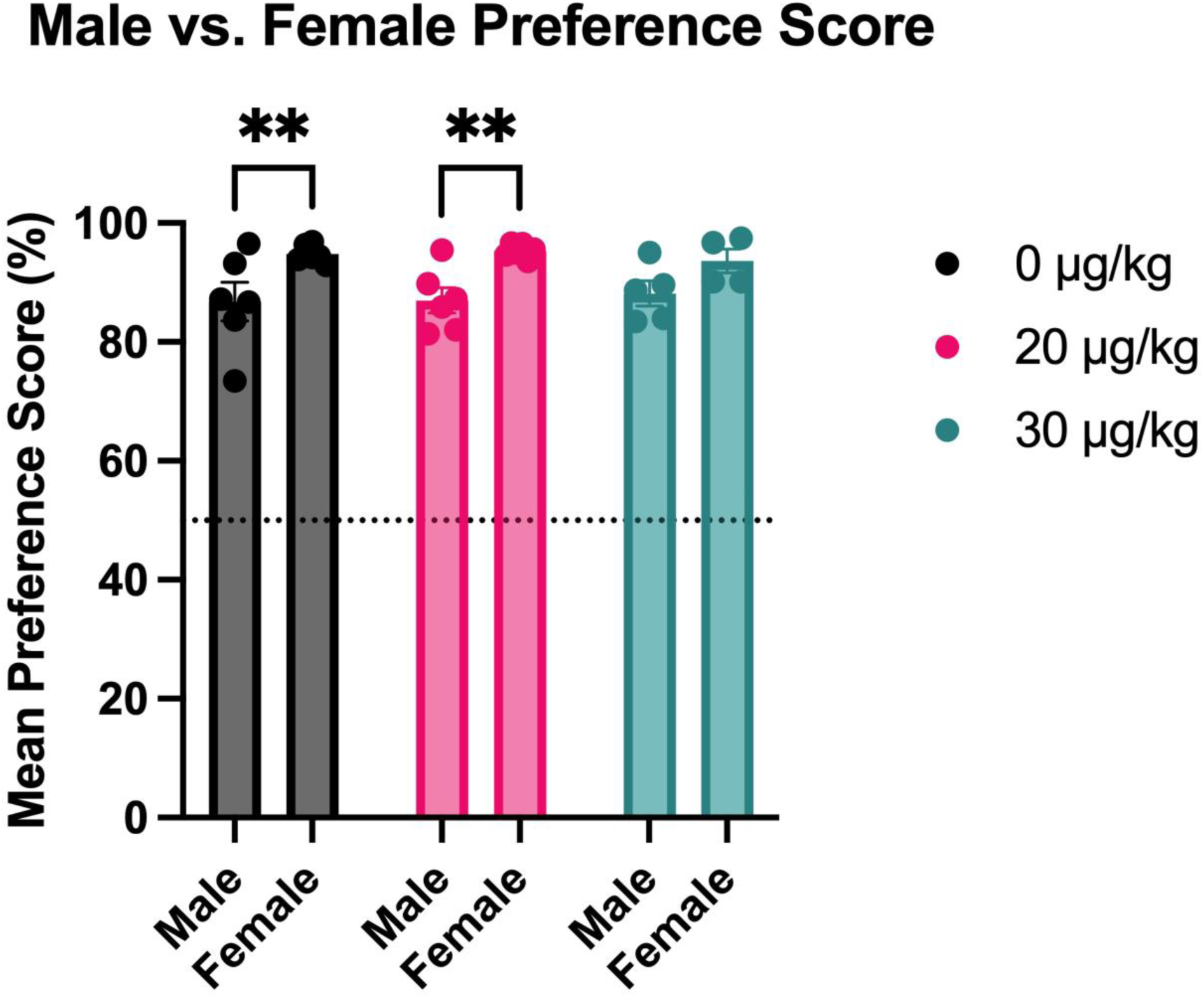
**Experiment 3: OVX females have a higher SACC preference than CAST males**. While there was no effect of G1 treatment, gonadectomy did reveal a basal sex difference between male and female SACC preference. OVX females in the 0 and 20 µg/kg treatment groups had a higher SACC preference than CAST males in the same treatment group (Effect of Sex: *p* < .001). Data of individual animals are shown as mean ± SEM. *p < 0.05, **p < 0.01.

### Fluid Intake

There were no significant treatment group differences in mean fluid intake in males or females. Fluid intake fluctuated significantly across days for males (Fig. 8B; Effect of Time: F_(3.154, 40.11)_ = 17.89, 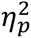 = 0.585) and females (Fig. 8D; Effect of Time: F_(2.920, 27.95)_ = 4.216, 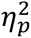 = 0.306)

**Figure 8.**
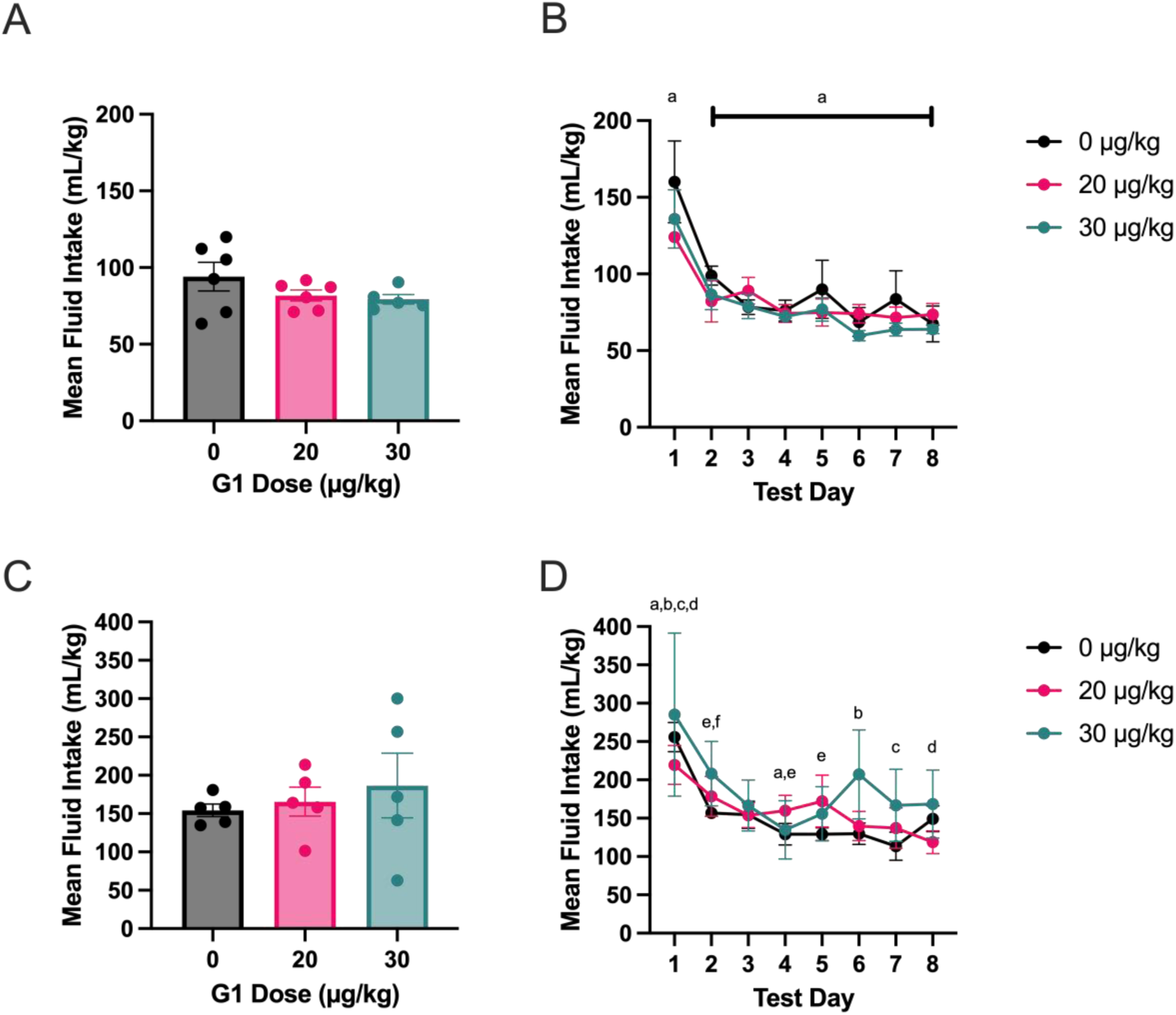
Experiment 3: Systemic administration of G1 does not affect fluid intake in gonadectomized males and females. (A&B) Males (C&D) Females. All groups showed similar patterns of fluid intake. Fluid intake fluctuated across days (Effect of Time: *p < .001*). (A&C) Data of individual animals are shown as mean ± SEM. (B&D) Group averages are shown as mean ± SEM. *Note. Days with different letters differ significantly (p < 0.05) as determined by Tukey post hoc tests*.

Once again, there were significant sex differences in mean fluid intake, with females consuming more than males (Fig. 9; Effect of Sex: F_(1,26)_ = 31.63, 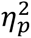 = .549). Males in all treatment groups had lower mean fluid intake than females in the same treatment group (Fig. 9; *d*’s = 2.873, 2.905, 1.609).

**Figure 9.**
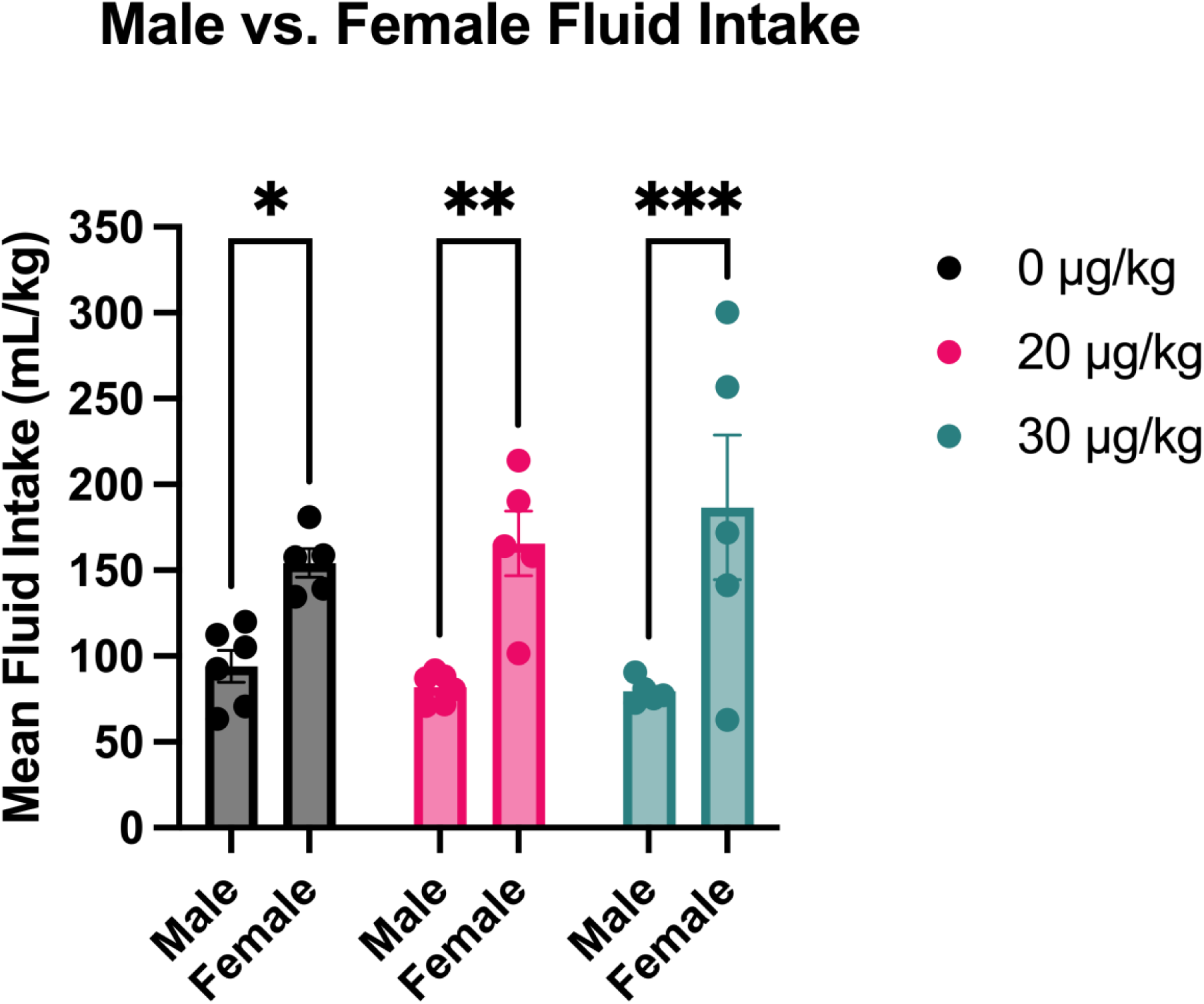
Experiment 3: OVX females drank more fluid than CAST males. While there was no effect of G1 treatment, gonadectomy did reveal a basal sex difference between male and female fluid intake (Effect of Sex: *p* < .001). OVX females drank more fluid than CAST males. Data of individual animals are shown as mean ± SEM. *p < 0.05, **p < 0.01, ***p < 0.001.

### SACC Intake

There were no significant treatment group differences in mean SACC intake in males. However, fluid intake did fluctuate significantly across days (Data Not Shown; Effect of Time: F_(2.755, 35.02)_ = 8.047, 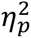 = 0.388). In females, there were no significant treatment group differences in mean SACC intake or in SACC intake across days.

Male vs. female mean SACC intake is shown in Table 3. Sex significantly impacted group differences in mean SACC intake between males and females (Table 3; Effect of Sex: F_(1,26)_ = 41.09, 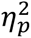 = 0.612). Overall, males had lower mean SACC intake than females regardless of treatment group (*d*’s = 2.84, 6.506, 1.635).

**Table 3.** Statistical analyses for Experiment 3: Mean SACC and Water Intake. Table shows data from a two-way ANOVA analysis of male and female mean SACC & water intake. There was a significant main effect of Sex on SACC intake. All male treatment groups drank significantly less SACC than their female counterparts. There were no group differences in water intake. *Note. S = sex; T = Treatment. Means with different subscripts differ with p < 0.05 as determined by Tukey post hoc tests. ****p < 0.0001*.

| Experiment 3: SACC & Water Intake |  |  |  |  |  |  |  |  |  |  |
| --- | --- | --- | --- | --- | --- | --- | --- | --- | --- | --- |
| Variable | 0 µg/kg G1 |  | 20 µg/kg |  | 30 µg/kg |  | Two-Way ANOVA |  |  |  |
| | *M* | *SD* | *M* | *SD* | *M* | *SD* | Effect | *F* | dF | $\eta_p^2$ |
| SACC Intake |  |  |  |  |  |  | S x T | 0.8103 | 2,26 | .059 |
| Male | 83.442<sup>a</sup> | 24.558 | 72.455<sup>b</sup> | 10.652 | 72.764<sup>c</sup> | 8.358 | S | 31.70<sup>****</sup> | 1,26 | 0.549 |
| Female | 143.360<sup>a</sup> | 13.349 | 158.714<sup>b</sup> | 40.889 | 179.472<sup>c</sup> | 93.569 | T | 0.268 | 2,26 | .020 |
| Water Intake |  |  |  |  |  |  | S x T | 0.047 | 2,26 | 0.004 |
| Male | 10.621 | 7.261 | 9.303 | 2.940 | 8.841 | 4.103 | S | 2.618 | 1,26 | 0.091 |
| Female | 7.861 | 2.112 | 6.903 | 2.115 | 7.139 | 0.783 | T | 0.323 | 2,26 | 0.024 |

### Water Intake

Mean water intake did not differ between treatment groups in males or females. However, water intake did fluctuate significantly across days in males (Data Not Shown; F_(4.002, 50.88)_ = 3.686, 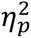 = 0.225) and females (Data Now Shown; F_(2.920, 27.95)_ = 4.216, 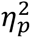 = 0.306).

There were no significant treatment group differences in water intake between males and females (Table 3).

### Summary

Systemic G1 administration did not significantly alter SACC preference or intake in either sex, despite fluctuations in fluid consumption over time.

Experiment 4: Systemic administration of G1 does not rapidly modulate 0.1% SACC preference in gonadectomized animals of either sex

### SACC Preference

Males and females in different treatment groups did not differ in mean preference SACC score (Fig. 10). However, preference scores did significantly differ across days in males (Fig. 10B; Effect of Time: F_(1.758, 22.42)_ = 64.66; 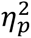 = 0.835) and females (Fig. 10D; Effect of Time: F_(2.256, 33.84)_ = 64.47, 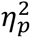 = 0.811) On day 2, males in the 0 µg/kg G1 group preferred SACC more than those in the 20 µg/kg group (*d* = 1.686). However, males and females preferred SACC at similar rates (Fig. 11).

**Figure 10.**
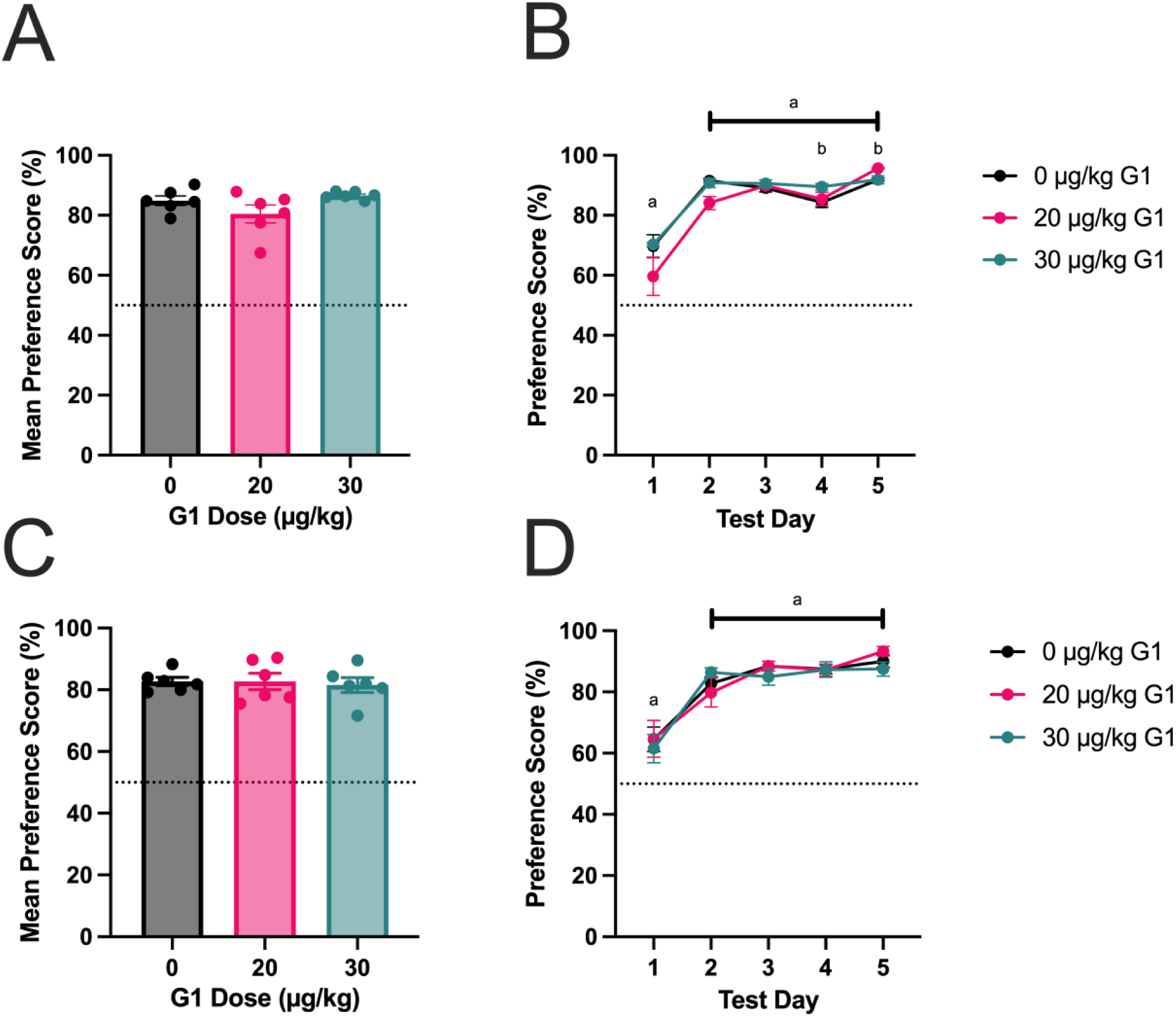
Experiment 4: G1 does not rapidly modulate SACC preference in gonadectomized males or females. (A&B) Males (C&D) Females. All treatment groups formed a SACC preference, as indicated by a mean preference score > 50%. Preference score fluctuated over time (Effect of time: *p < .001*). (A&C) Data of individual animals are shown as mean ± SEM. (B&D) Group averages are shown as mean ± SEM. *Note. Days with different letters differ significantly (p < 0.05) as determined by Tukey post hoc tests*.

**Figure 11.**
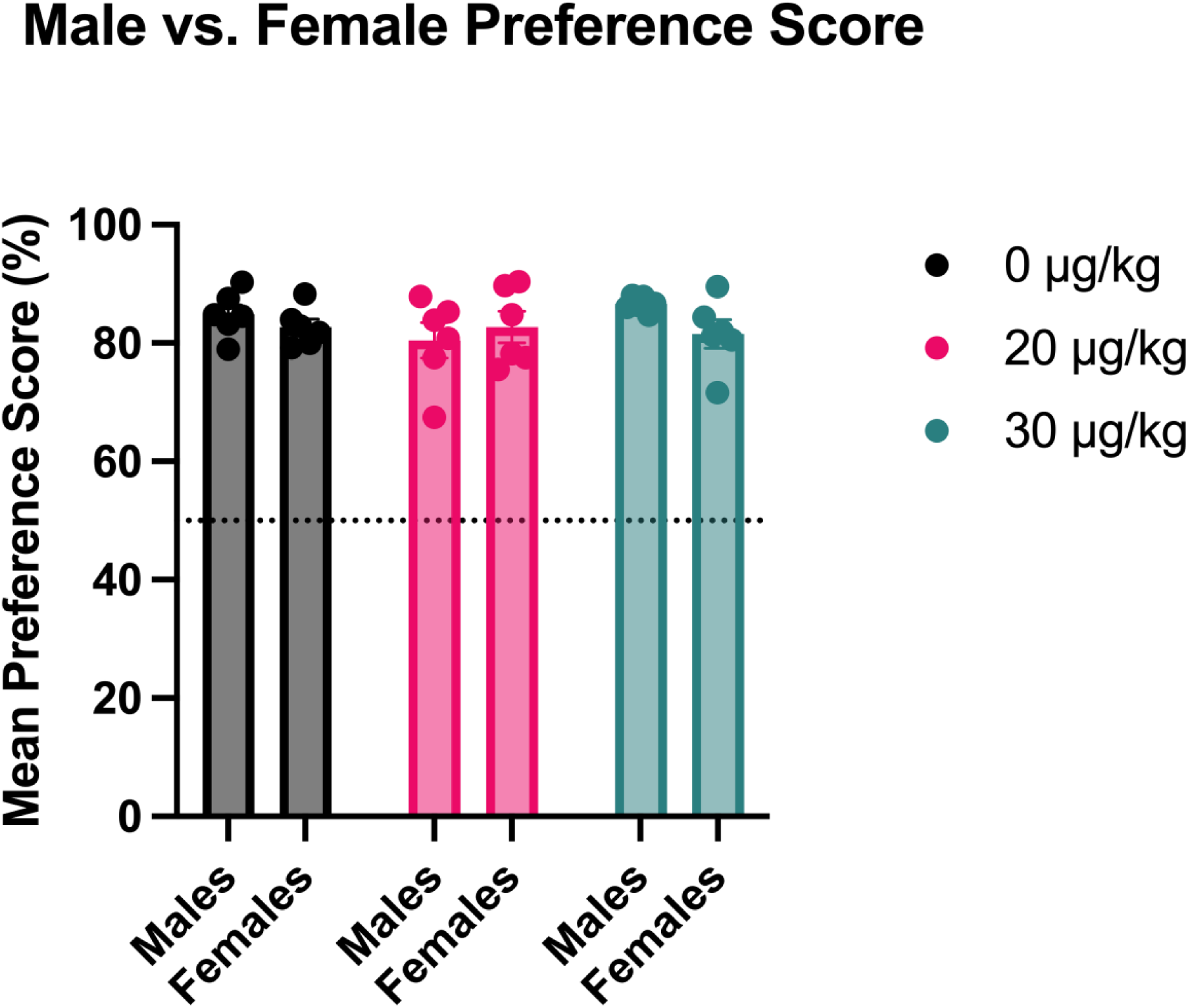
Experiment 4: SACC preference did not differ between sexes. Males and females preferred SACC at similar rates. Data of individual animals are shown as mean ± SEM.

### Fluid Intake

Both males and females across treatment groups consumed similar amounts of fluid overall. However, the amount consumed varied over time in males (Fig. 12B; Effect of Time: F_(3.165, 40.36)_ = 4.685, 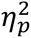 = 0.269) and females (Fig. 12D; Effect of Time: F_(2.600, 39.00)_ = 8.648, 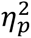 = 0.366). In males, fluid intake was greater on day five than day one (*d* =1.943). In females, fluid intake was lower on day one compared to day four (*d* = 5.927) and five (*d* = 2.448) and lower on day two compared to day four (*d* =15.001) and five (*d* = 3.698). Males and females did not drink different amounts of fluid.

**Figure 12.**
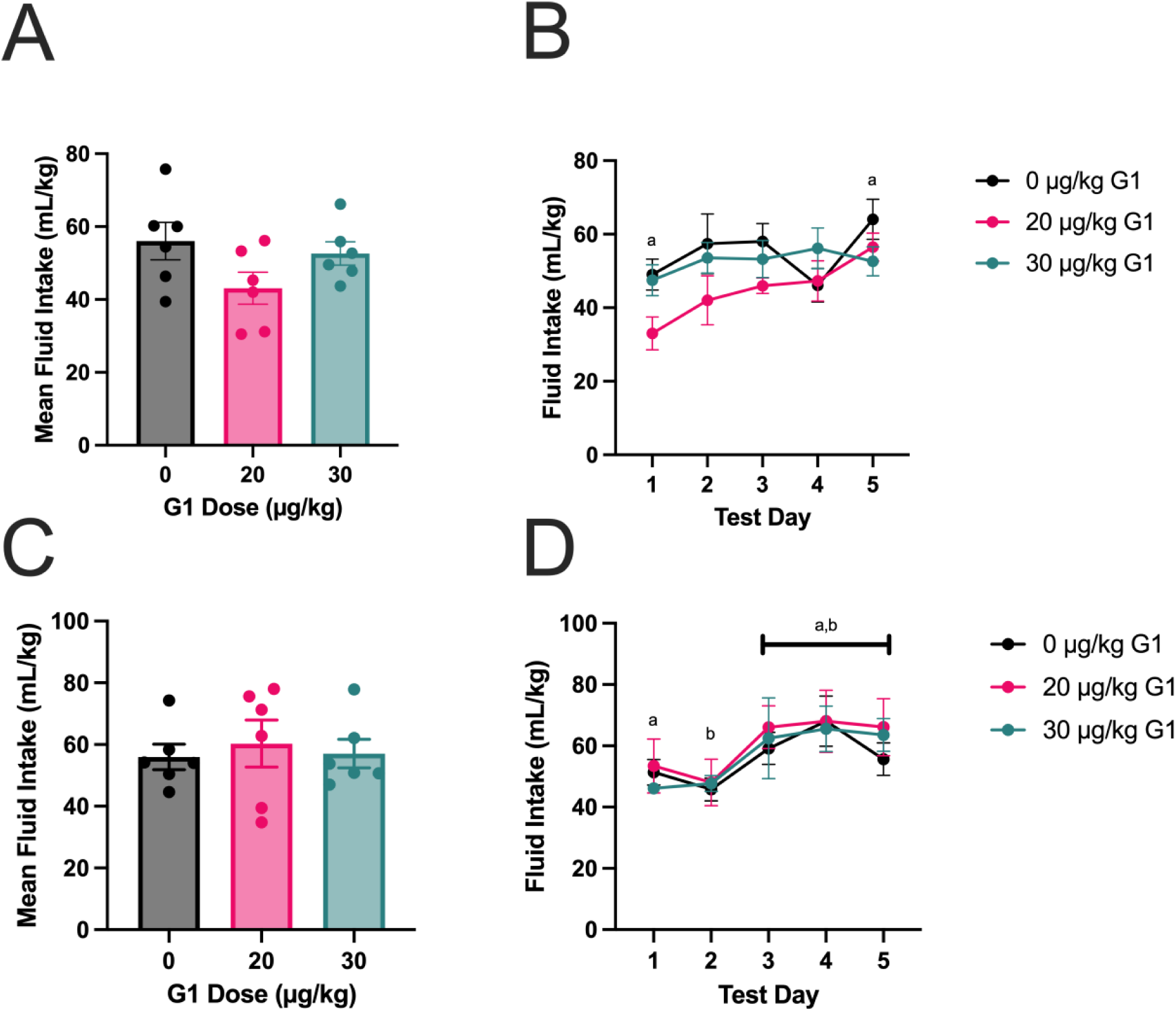
Experiment 4: G1 does not rapidly modulate fluid intake in gonadectomized animals. (A&B) Males (C&D) Females. All treatment groups drank similar amounts of fluid. Fluid intake fluctuated over time (Effect of time: *p < .001*). Data of individual animals are shown as mean ± SEM. (A&C) Data of individual animals are shown as mean ± SEM. (B&D) Group averages are shown as mean ± SEM. *Note. Days with different letters differ significantly (p < 0.05) as determined by Tukey post hoc tests*.

**Figure 13.**
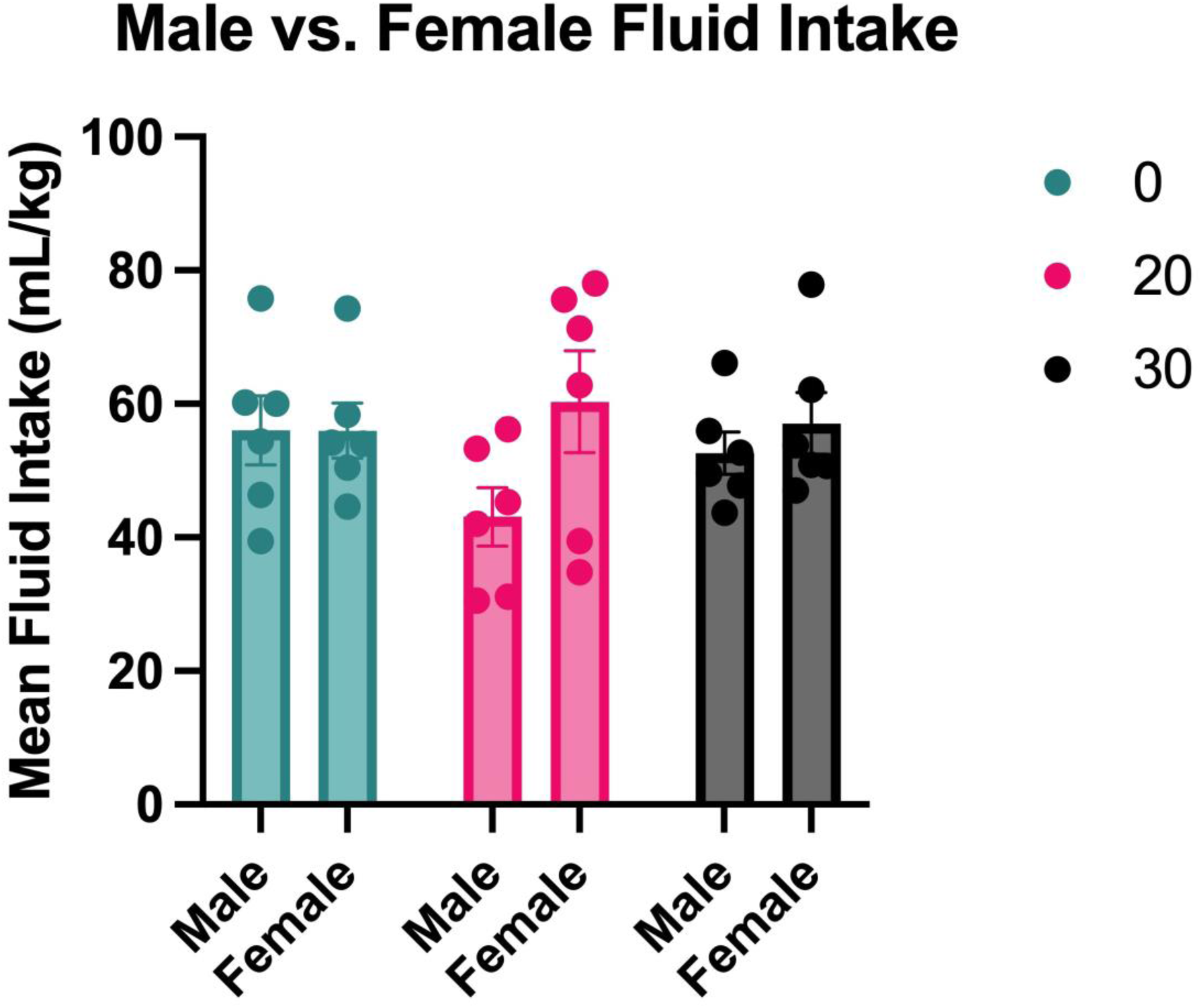
Experiment 4: No sex differences in mean fluid intake. Males and females consumed similar amounts of fluid. Data of individual animals are shown as mean ± SEM.

### SACC Intake

Male and female mean SACC intake data can be found in Table 4. Different treatment groups did not differ in their mean SACC intake within sex. However, SACC intake fluctuated across days for both males (Table 4; Effect of Time: F_(3.291, 41.97)_ = 18.63, 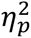 = 0.594) and females (Table 4; Effect of Time: F_(2.553, 38.29)_ = 19.26, 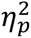 = 0.562). In males, SACC intake on day one was lower than on all subsequent days (*d =* 1.931, 2.629, 1.976, 3.664). In females, SACC intake on day one was lower than on all subsequent days (*d =* 2.495, 6.394, 9.545, 4.754). Additionally, in females, there were significant differences between day two and day three (*d =* 6.039), four (*d =* 14.833), and five (*d =* 3.790). Males and females drank similar amounts of SACC.

**Table 4.**
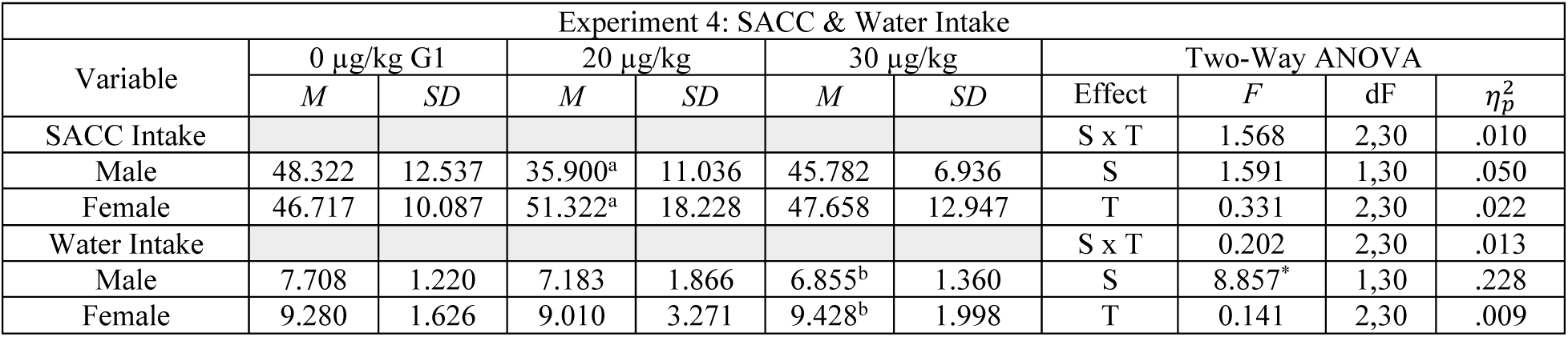
Statistical analyses for Experiment 4: Mean SACC and Water Intake. Table shows data from a two-way ANOVA analysis of male and female mean SACC & water intake. There were no group differences in SACC intake. There was a significant effect of Sex on water intake. Females in the 30% treatment group drank more water than males in the same group. *Note. S = sex; T = treatment. Means with different subscripts differ with p < 0.05 as determined by Tukey post hoc tests. *p < 0.05*.

### Water Intake

Male and female mean water intake data can be found in Table 4. Males in different treatment groups did not differ in their mean water intake. However, differences in water intake across days emerged (Table 5; Effect of Time: F_(2.200, 28.05)_ = 49.01, 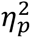 = 0.794). On day 2, males in the 20 µg/kg G1 group drank less water than those in other treatment groups (*p* < .0001 & *p* = 0.0326). Females in different treatment groups did not differ in their mean water intake.

**Table 5.** Statistical analyses for Experiment 4: Male Water Intake. Table shows data from a mixed-effect analysis of male water intake across days. *Note: Days with a superscript were significantly different (p < 0.05) as determined by Tukey post hoc tests.* *^a^ Significantly different from D1* >*^b^ Significantly different from D4*

| Experiment 4: Male Water Intake |  |  |  |  |  |  |
| --- | --- | --- | --- | --- | --- | --- |
| Variable | 0 µg/kg G1 |  | 10 µg/kg G1 |  | 20 µg/kg G1 |  |
| *Water Intake* | *M* | *SD* | *M* | *SD* | *M* | *SD* |
| D1 | 14.391 | 4.336 | 12.659 | 4.221 | 14.213 | 3.701 |
| D2<sup>a</sup> | 4.535 | 1.230 | 6.302 | 2.349 | 5.068 | 2.583 |
| D3<sup>a</sup> | 6.133 | 1.244 | 4.588 | 0.393 | 5.231 | 2.694 |
| D4<sup>a</sup> | 7.029 | 0.705 | 6.598 | 2.546 | 5.710 | 1.158 |
| D5<sup>a,b</sup> | 5.109 | 0.210 | 2.397 | 0.195 | 4.045 | 1.114 |

However, water intake fluctuated across days (Data Not Shown; Effect of Time: F_(1.858, 27.87)_ = 36.31; 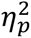 = 0.708). Males and females drank different amounts of water (Table 4; Effect of Sex: F_(1, 30)_ = 8.857; 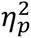 = 0.228).

### Summary

Systemic G1 administration did not rapidly alter SACC preference or intake in either sex, despite fluctuations in fluid consumption across days.

Experiment 5: Chronic systemic administration of G1 does not modulate 0.1% SACC preference in intact animals of either sex

### SACC Preference

Males in both treatment groups showed similar SACC preference rates, with no significant differences between treatment conditions (Fig. 14A&B). Similarly, females in both treatment conditions preferred SACC at comparable rates (Fig. 14C&D). Overall, there were no sex differences in SACC preference (Fig. 15).

**Figure 14.**
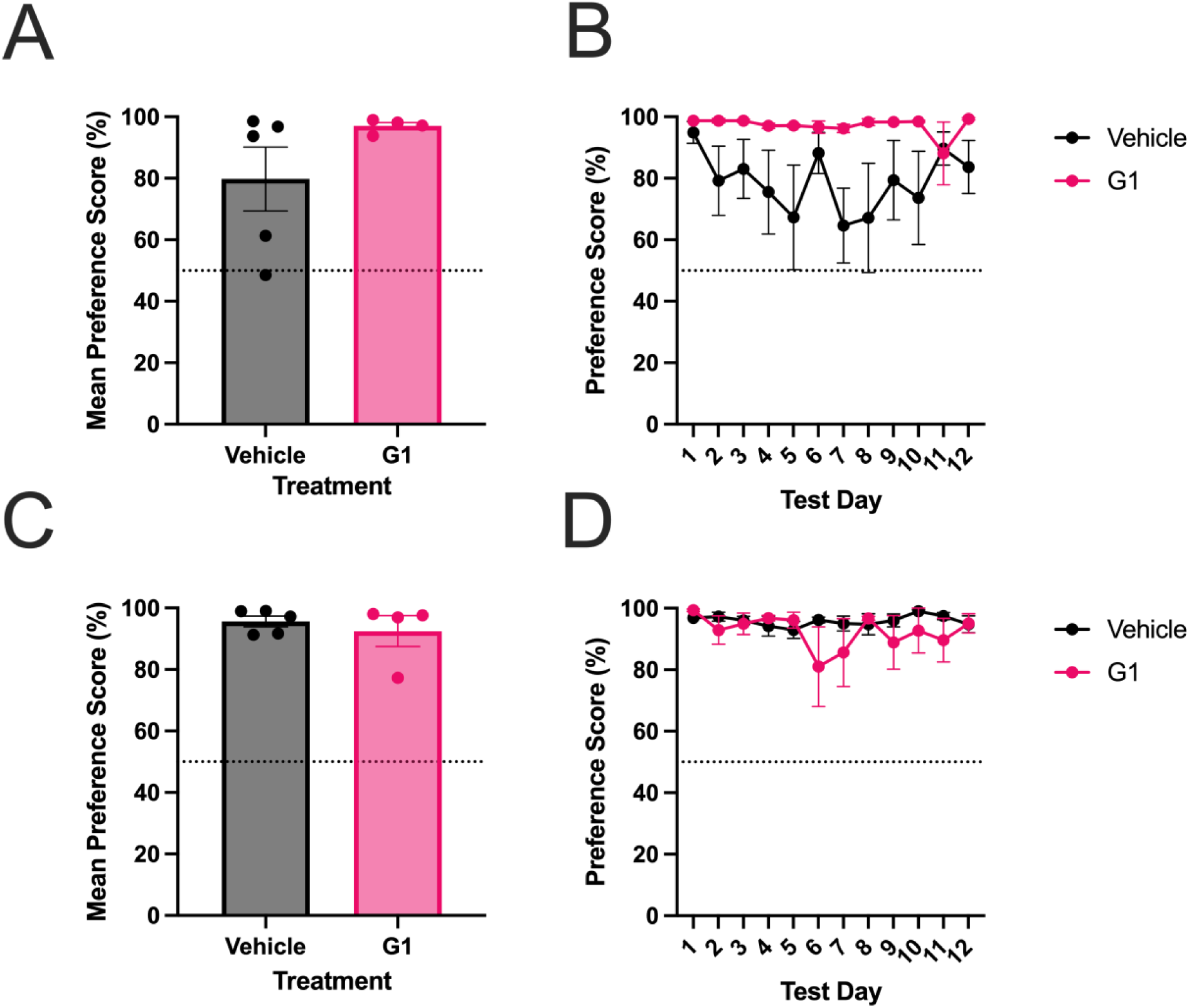
Experiment 5: Chronic G1 administration does not affect preference score in males or females. (A&B) Males (C&D) Females. Chronic administration of G1 did not affect mean preference score or preference score across days. (A&C) Data of individual animals are shown as mean ± SEM. (B&D) Group averages are shown as mean ± SEM.

**Figure 15.**
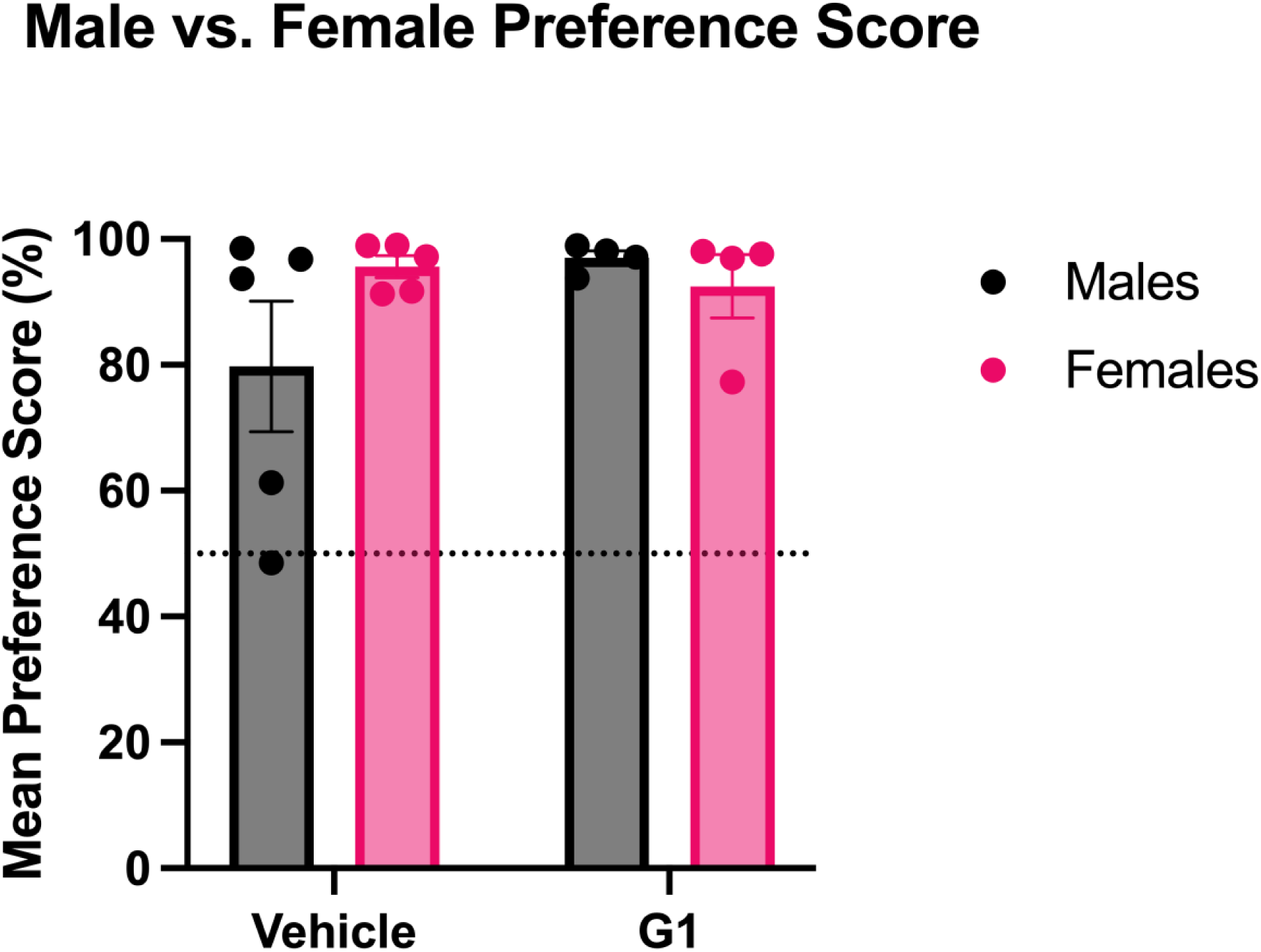
Experiment 5: SACC preference did not differ between sexes. There were no sex differences in mean preference scores between males and females. Data from individual animals are shown as mean ± SEM.

### Fluid Intake

Overall, males in both treatment groups consumed similar mean amounts of fluid. Males had differences in fluid intake across days (Fig. 16B; Main Effect of Time: F_(3.540, 22.21)_ = 4.429, 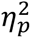 = 0.416), however post hoc analyses did not reveal any group differences.

**Figure 16.**
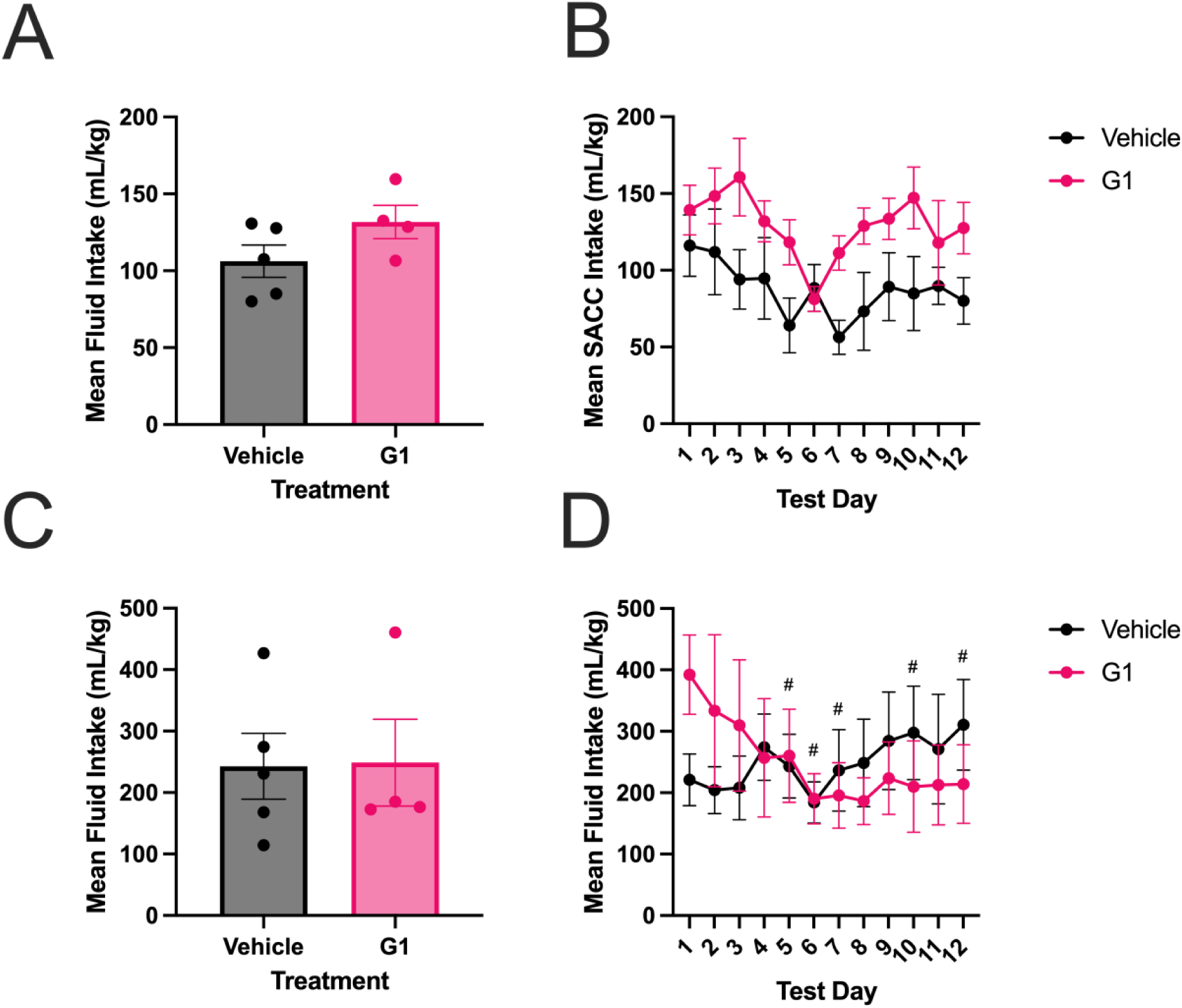
Experiment 5: Chronic G1 does not modulate mean fluid intake. (A&C) Males (C&D) Females. Both sexes’ treatment groups consumed the same mean amount of fluid. However, G1-treated female fluid intake fluctuated across days. (A&C) Data of individual animals are shown as mean ± SEM. (B&D) Group averages are shown as mean ± SEM. *Note. In G1-treated females, days with # differ significantly from day one (p < 0.05) as determined by Tukey post hoc tests*.

**Figure 17.**
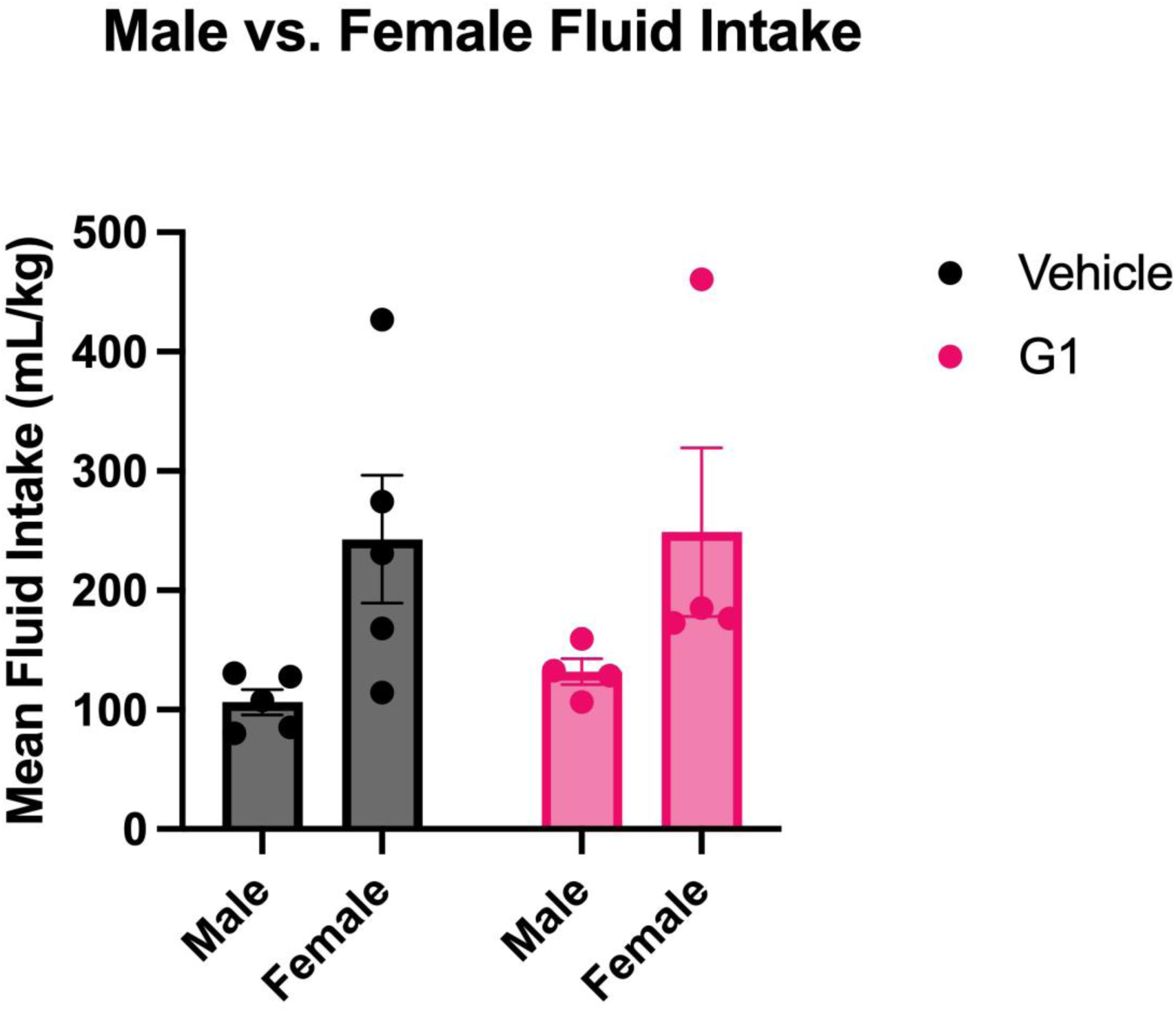
Experiment 5: No sex differences in mean fluid intake. Mean fluid intake did not differ between sexes in either treatment group. Data of individual animals are shown as mean ± SEM. **p < 0.01

In females, there was a Time x Treatment interaction (Fig. 16D; F_(11,73)_ = 5.037, 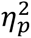 = 0.431). In G1-treated females fluid intake on day one differed from day five (*d =* 0.939), six (*d* = 1.871), seven (*d* = 1.662), ten (*d* = 1.311), and twelve (*d* = 1.387).

Furthermore, there was a significant difference in mean fluid intake between males and females (Fig. 16; Effect of Sex: F_(1, 14)_ = 8.296; 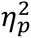 = 0.372).

### SACC Intake

Both males and females drank similar amounts of SACC across treatment conditions (Table 6). In males, there were differences in SACC intake across days (Table 7; Main Effect of Time: F_(3.419, 21.45)_ = 4.545; 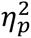 = 0.420). SACC intake in vehicle-treated males was significantly lower than that of G1-treated males on day 7 (*d* = 2.463).

**Table 6.** Statistical analyses for Experiment 5: Mean SACC and Water Intake. Table shows data from a two-way ANOVA analysis of male and female mean SACC & water intake. There was a significant main effect of Treatment with 0 µg/kg males drinking significantly less SACC than their female counterparts. There were no group differences in water intake. *Note. S = sex; T = treatment. *p < 0.05*.

| Experiment 5: SACC & Water Intake |  |  |  |  |  |  |  |  |
| --- | --- | --- | --- | --- | --- | --- | --- | --- |
| Variable | 0 $\mu\text{g/kg}$ G1 | | 90 $\mu\text{g/kg}$ | | Two-Way ANOVA | | | |
| | *M* | *SD* | *M* | *SD* | Effect | *F* | dF | $\eta_p^2$ |
| SACC Intake | | | | | S × T | 0.188 | 1,14 | .013 |
| Male | 88.902 | 41.104 | 128.650 | 23.391 | S* | 7.653 | 1,14 | .353 |
| Female | 235.640 | 122.653 | 236.375 | 144.990 | T | 0.194 | 1,14 | .014 |
| Water Intake | | | | | S × T | 2.884 | 1,14 | .171 |
| Male | 17.377 | 18.626 | 3.121 | 1.691 | S | 0.005 | 1,14 | .000 |
| Female | 7.255 | 4.623 | 12.425 | 13.587 | T | 0.631 | 1,14 | .043 |

**Table 7.** Statistical analyses for Experiment 5: SACC and Water Intake. Table shows data from two-way ANOVA analyses of male and female SACC & water intake across days. *Note: Days with a superscript were significantly different (p < 0.05) as determined by Tukey post hoc tests*.

| Experiment 5: Male and Female SACC and Water Intake Across Days |  |  |  |  |  |  |  |  |
| --- | --- | --- | --- | --- | --- | --- | --- | --- |
| Variable | Male |  |  |  | Female |  |  |  |
|  | Vehicle |  | G1 |  | Vehicle |  | G1 |  |
|  | *M* | *SD* | *M* | *SD* | *M* | *SD* | *M* | *SD* |
| *SACC Intake* |  |  |  |  |  |  |  |  |
| D1 | 116.106 | 44.613 | 139.287 | 27.997 | 215.430 | 96.289 | 389.255<sup>abc</sup> | 124.967 |
| D2 | 111.966 | 62.491 | 148.433 | 36.299 | 200.138 | 87.202 | 318.323 | 257.937 |
| D3 | 94.050 | 43.260 | 160.675 | 50.391 | 201.180 | 104.462 | 296.703 | 210.925 |
| D4 | 94.710 | 59.169 | 131.887 | 23.021 | 264.810 | 129.641 | 249.648 | 188.405 |
| D5 | 64.043 | 35.688 | 118.260 | 29.220 | 231.544 | 120.933 | 250.715<sup>a</sup> | 150.449 |
| D6 | 88.495 | 30.503 | 81.287 | 13.949 | 177.846 | 74.288 | 160.735 | 98.049 |
| D7 | 56.358 | 22.273 | 111.298 | 22.334 | 229.682 | 153.969 | 174.100<sup>b</sup> | 118.908 |
| D8 | 73.288 | 50.726 | 128.898 | 23.497 | 239.498 | 146.988 | 180.445 | 74.621 |
| D9 | 89.328 | 49.412 | 133.500 | 26.992 | 277.754 | 180.305 | 208.378 | 131.290 |
| D10 | 84.878 | 53.853 | 147.230 | 34.780 | 294.673 | 150.663 | 201.990 | 156.239 |
| D11 | 89.814 | 27.119 | 117.965 | 55.009 | 266.463 | 178.915 | 199.305 | 139.231 |
| D12 | 80.084 | 33.989 | 127.513 | 33.628 | 301.398 | 170.808 | 206.860<sup>c</sup> | 131.037 |
| *Water Intake* |  |  |  |  |  |  |  |  |
| D1 | 4.338 | 6.099 | 1.617 | 1.402 | 5.500 | 3.694 | 3.213 | 4.334 |
| D2 | 19.228 | 17.998 | 1.790 | 2.250 | 4.058 | 3.967 | 15.233 | 20.369 |
| D3 | 13.878 | 16.611 | 1.808 | 1.208 | 6.828 | 3.889 | 12.900 | 13.670 |
| D4 | 21.858 | 25.857 | 3.807 | 1.432 | 9.298 | 10.150 | 7.168 | 4.621 |
| D5 | 27.818 | 26.968 | 3.443 | 1.382 | 11.746 | 6.318 | 9.530 | 11.175 |
| D6 | 9.800 | 9.459 | 2.900 | 3.365 | 6.342 | 2.673 | 29.458 | 34.543 |
| D7 | 30.625 | 21.140 | 4.038 | 2.416 | 6.780 | 6.566 | 21.518 | 29.563 |
| D8 | 27.820 | 29.546 | 2.205 | 3.074 | 9.013 | 10.600 | 5.905 | 2.585 |
| D9 | 19.580 | 28.054 | 2.208 | 0.106 | 6.816 | 6.384 | 15.368 | 20.338 |
| D10 | 21.276 | 26.208 | 2.157 | 0.117 | 2.853 | 1.946 | 8.008 | 16.015 |
| D11 | 8.918 | 10.089 | 9.505 | 14.751 | 4.568 | 1.620 | 13.528 | 14.551 |
| D12 | 12.852 | 15.065 | 1.033 | 1.192 | 9.160 | 7.603 | 7.285 | 6.806 |

In females, there was a Time x Treatment interaction (Table 7; F_(11,73)_ = 4.770; 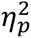 = 0.418). Males and females drank different amounts of SACC (Table 6; Effect of Treatment: F_(1, 12)_ = 8.351; 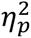 = 0.410). Specifically, vehicle-treated females drank more SACC than vehicle-treated males (*d* = 1.603). G1-treated female SACC intake on day one differed from day five (*d* = 1.543), seven (*d* = 1.763), and twelve (1.495).

### Water Intake

Males in both treatment groups drank similar amounts of water across days. There was a Time x Treatment interaction in females (Table 7; F_(11, 73)_ = 1.993; 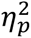 = 0.231). Post hoc analyses did not reveal any group differences. There were no sex differences in water intake (Table 7).

### Summary

Chronic administration of G1 did not significantly alter SACC preference in either sex.

## General Discussion

These studies demonstrate that GPER-1 activation modulates reward preference across different reward types (saccharin vs. cocaine) in a sex- and site-dependent manner. Using both intra-DLS and systemic administration of G1, we examined its effects on SACC preference and cocaine CPP across different timescales. We found that intra-DLS GPER-1 activity attenuates SACC preference in males, but not females, in a dose-dependent fashion (see Fig. 1A&C). Systemic administration of G1 blocked cocaine-induced CPP in both sexes (see Fig. 5). However, systemic administration of G1 did not change SACC preference in either sex under any experimental conditions. These findings suggest that GPER-1 effects are sex-, site-, and reward-dependent.

### Intra-DLS G1 Saccharin Preference Studies

Experiment 1 revealed that 20% G1 intra-DLS administration significantly reduced SACC preference in males but did not affect females. Males in this treatment group showed a mean SACC preference of 39% (Fig. 1A), suggesting that intra-DLS GPER-1 activation modulated SACC consummatory behavior. Determining how intra-DLS GPER-1 activation alters SACC preference in males will require further investigation.

One possible explanation for this result involves modulation of the valence of SACC. Rats generally prefer saccharin, and while the taste of saccharin can have mildly aversive components (Dess, 1993; Kuhn et al., 2004), the 0.1% concentration used here falls below the threshold for being aversive in rats (Bachmanov et al., 2001; Rehn et al., 2018; Sclafani et al., 2010). This suggests that intra-DLS GPER-1 activation either lowered the hedonic value or increased the aversive qualities of saccharin. In support of the hedonic value theory, one study found that male rats administered tamoxifen (a SERM and GPER-1 agonist) developed a conditioned taste avoidance of sucrose during a single bottle lickometer test but not an increase in active rejection reactions. In support of the aversion theory, systemic administration of G1 has been shown to regulate aversively motivated memory in male rats (de Souza et al., 2021). G1 treatment enhances inhibitory avoidance when administered immediately following training, likely through emotional memory formation mechanisms (de Souza et al., 2021). GPER-1 activation can influence behavior and memory, enhancing the salience of negative or aversive memories. This raises the possibility that intra-DLS GPER-1 activation strengthened the encoding of negative or aversive aspects of the SACC experience, leading to a reduced preference.

Another potential mechanism involves differences in fluid balance regulation. GPER-1 has been implicated in regulating fluid balance (Qiao et al., 2016; Santollo et al., 2013; Santollo and Daniels, 2015). Previous studies have observed that intracerebroventricular (ICV) infusions of G1 decrease angiotensin II-stimulated water intake in ovariectomized females (Santollo and Daniels, 2015). However, in the present study, intra-DLS GPER-1 activation did not affect overall fluid intake in males, suggesting that intra-DLS GPER-1 activity does not regulate fluid balance in males. While sex-specific effects were observed–males in the 20 and 30% G1-treated groups consumed less total fluid than their female counterparts–this difference was likely driven primarily by a shift in preference rather than a general suppression of fluid intake. Specifically, males drank more water and less SACC, reinforcing the idea that intra-DLS G1 administration altered the reward valence of SACC rather than influencing fluid homeostasis through changes in thirst.

Further analysis of drinking microstructure (e.g., lick rates, burst patterns) and taste reactivity (e.g., hedonic facial expressions) will be critical for determining whether intra-DLS GPER-1 activation alters SACC preference through changes in palatability, associative learning, or reward valuation.

### Conditioned Place Preference

Past studies showed that GPER-1 signaling modulates the development of a drug-induced conditioned place preference. Specifically, intra-DLS activation of GPER-1 attenuates cocaine-induced CPP in males, while morphine-induced CPP is potentiated in GPER-1 KO males (Quigley and Becker, 2021; Sun et al., 2020). Intra-DLS G1 administration did not affect cocaine-induced CPP in intact females (citation).

The current results expand on those findings by demonstrating that systemic administration of 20 µg/kg G1 during conditioning sessions not only blocks the development of a 10 mg/kg cocaine-induced CPP in both males and females (Fig. 5). This result was particularly surprising given that (a) intra-DLS G1 administration did not affect SACC preference or block a cocaine-induced CPP in females, and (b) intra-DLS G1 administration has been shown to enhance motivation for cocaine in females (citation). These discrepancies suggest that systemic and intra-DLS administration of G1 engage distinct neural mechanisms in males and females.

One possible explanation is that intra-DLS GPER-1 activation selectively influences reward-related sensory processing in males but not females. Cocaine’s reinforcing effects are entirely pharmacological, directly inducing synaptic dopamine overflow. In contrast, saccharin engages both sensory and reward-processing systems. Suppose intra-DLS GPER-1 activity modulates with sensory aspects of reward processing in males but not females. In that case, it may explain why intra-DLS G1 reduced SACC preference in males but had no effect in females, while systemic G1 blocked cocaine CPP in both sexes.

Alternatively, the results of the CPP experiment could also be explained by an aversive state induced by G1 administration itself, reducing cocaine reward rather than directly blocking GPER-1 mediated reinforcement mechanisms. While animals did not significantly shift their preference to the saline-paired chamber (Fig. 5), the experimental design involved G1 administration prior to drug- **and** saline-conditioning sessions. Future studies should restrict G1 administration to drug conditioning sessions to assess whether a stronger preference for the saline-paired side develops. If not, this further supports the theory that systemic G1 may have diminished the rewarding effects of cocaine.

Finally, in male mice, GPER1 knockout increased sensitivity to morphine-induced CPP but it did not alter sensitivity to sucrose-induced CPP (Sun et al., 2020). These data suggest that GPER-1 regulation of reward processing depends on both the site of activation and the reward being studied. The current findings further support this idea, as intra-DLS G1 administration selectively altered SACC preference in males, while systemic G1 blocked cocaine CPP in both sexes. Understanding the precise mechanisms by which GPER-1 modulates reinforcement and reward valence will require further investigation into its interactions with dopaminergic and sensory processing circuits.

### Systemic G1 Saccharin Preference Studies

Systemic G1 administration did not alter SACC preference under any experimental condition, regardless of treatment duration, sex, or gonadal status. Neither acute nor chronic G1 administration affected SACC preference or intake in gonadectomized animals, males or females. This finding contrasts with the intra-DLS G1 administration, which selectively reduced SACC preference in intact males, further highlighting the importance of the site of action and a potential role for circulating gonadal hormones.

Interestingly, chronic systemic (but not intra-DLS) G1 administration in intact females resulted in time-dependent modulation of fluid intake. However, this effect was non-selective and did not correspond with changes in SACC preference. The data suggests that GPER-1 signaling may contribute to fluid homeostasis in a manner distinct from its role in reward processing. It was previously mentioned that Santollo & Daniels (2015) found that ICV administration of G1 reduced angiotensin II-induced fluid intake in OVX females. GPER-1, also an aldosterone target, is expressed in brain regions like the paraventricular nucleus, supraoptic nucleus, and nucleus tractus solitarius, all vital to fluid homeostasis (Brailoiu et al., 2007; Qiao et al., 2016; Sakamoto et al., 2007; Santollo and Daniels, 2015). Additionally, GPER-1 signaling may reduce aldosterone receptor expression levels, which may mediate attenuated angiotensin II-induced fluid intake (Koganti et al., 2014).

The lack of an effect of systemic G1 on SACC preference suggests that GPER-1 signaling does not broadly regular reward-related behaviors but may have site-specific roles in modulating reinforcement and sensory processing. In males specifically, intra-GPER-1 activation selectively alters sensory or reward processing, while systemic GPER-1 signaling broadly modulates processes such as fluid balance without significantly engaging reward-related circuitry.

### Future Studies

Future studies will be conducted to address some of the limitations of the present experimental design. First, the studies simultaneously introduce SACC and G1 treatments. Several studies have demonstrated that GPER-1 signaling modulates learning and memory (De Souza et al., 2021; Gabor et al., 2015; Hammond et al., 2012, 2009; Lymer et al., 2017, 2018). The acquisition of a SACC preference may be dissociable from the sustained expression of said preference, and GPER-1 may modulate different components in males vs. females. Future studies should alter the timing of SACC and treatment introduction to understand learning processes better. Second, E2 levels regulate GPER-1 expression, and both subsequently change following gonadectomy (Cheng et al., 2014; Llorente et al., 2020; Marraudino et al., 2021; Spary et al., 2013). Animals in the acute systemic administration study were gonadectomized at least a week before G1 treatment. Whether or not these changes will impact the results remains to be seen. Replicating the experiment will increase confidence in the results.

## Conclusions

In conclusion, the present findings indicate that GPER-1’s effects are reward-, site-, and sex-specific. We confirmed that intra-DLS GPER-1 administration attenuates SACC preference in males but not females, likely due to sex differences in how GPER-1 modulates the integration of sensory information vs. reward reinforcement. In males, GPER-1 may influence the motivational aspects of reward more generally. Systemic G1 effectively reduces cocaine-induced CPP in both sexes, a surprising finding considering that intra-DLS administration only attenuated cocaine-induced CPP in males (Quigley and Becker, 2021), suggesting that intra-DLS vs systemic G1 affects cocaine reinforcement differently. These results highlight the potential of GPER-1 as a novel target for selectively modulating reward preferences in males and females.

## Notes

### Competing Interest Statement

The authors have declared no competing interest.

